# The Alternative Sigma Factor SigL Influences *Clostridioides difficile* Toxin Production, Sporulation, and Cell Surface Properties

**DOI:** 10.1101/2022.01.26.477884

**Authors:** Andrew E. Clark, Chelsea C. Adamson, Katelyn E. Carothers, Bryan Angelo P. Roxas, V.K. Viswanathan, Gayatri Vedantam

## Abstract

The alternative sigma factor SigL (Sigma-54) facilitates bacterial adaptation to the extracellular environment by modulating the expression of defined gene subsets. A homolog of the gene encoding SigL is conserved in the diarrheagenic pathogen *Clostridioides difficile*. To explore the contribution of SigL to *C. difficile* biology, we generated *sigL*-disruption mutants (*sigL::erm*) in strains belonging to two phylogenetically distinct lineages – the human-relevant Ribotype 027 (strain BI-1) and the veterinary-relevant Ribotype 078 (strain CDC1). Comparative proteomics analyses of mutants and isogenic parental strains revealed lineage-specific SigL regulons. Concomitantly, loss of SigL resulted in pleotropic and distinct phenotypic alterations in the two strains. Sporulation kinetics, biofilm formation, and cell surface-associated phenotypes were altered in CDC1 *sigL::erm* relative to the isogenic parent strain, but remained unchanged in BI-1 *sigL::erm*. In contrast, secreted toxin levels were significantly elevated only in the BI-1 *sigL::erm* mutant relative to its isogenic parent. We also engineered SigL overexpressing strains and observed enhanced biofilm formation in the CDC1 background, and reduced spore titers as well as dampened sporulation kinetics in both strains. Thus, we contend that SigL is a key, pleiotropic regulator that dynamically influences *C. difficile*’s virulence factor landscape, and thereby, its interactions with host tissues and co-resident microbes.

**Contribution to the Field:** Alternative sigma factors modulate bacterial gene expression in response to environmental and metabolic cues. *C. difficile* encodes multiple alternative sigma factors which mediate critical cellular processes. A Sigma-54 ortholog, SigL, is conserved across all sequenced strains, yet its function remains to be elucidated. We demonstrate that SigL dynamically influences *C. difficile* biology in fundamental, but distinct, ways in two strains representing two clinically-relevant *C. difficile* phylogenetic lineages (ribotypes). Loss of SigL results in changes in sporulation kinetics, biofilm formation, alterations in cell surface properties, and secreted toxin levels in a strain-specific manner. SigL overexpression, however, uniformly impedes sporulation. Thus, SigL plays a fundamental role in regulating the expression of *C. difficile* virulence factors and likely profoundly impacts pathogenesis in a strain-specific manner.

## Introduction

*Clostridioides difficile*, an anaerobic, spore-forming, Gram-positive bacillus, is a significant source of healthcare-associated infections (Stewart et al., 2020). *C. difficile* infection manifests as a diverse spectrum of diarrheal disease with possible progression toward more serious sequelae including pseudomembranous colitis or fatal toxic megacolon (George et al., 1978;Dudukgian et al., 2009). Ingested spores germinate into vegetative cells upon exposure to specific bile salt and amino acid triggers in the mammalian host. Disruption or suppression of the endogenous microbiota by antibiotics, immunosuppressive drugs, or host factors, facilitate vegetative cell colonization of the intestine. Vegetative cells from toxigenic *C. difficile* strains produce two major enterotoxins (TcdA and TcdB) that glucosylate host Rho GTPases, leading to actin cytoskeleton reorganization, solute transport disruption, and consequent symptomatic disease (Vedantam et al., 2012). *C. difficile* toxin synthesis increases as bacteria enter the stationary phase of growth; thus pathogen metabolism and virulence are linked (Voth and Ballard, 2005;Dineen et al., 2010;Bouillaut et al., 2015).

Sigma factors are dissociable components of the RNA polymerase holoenzyme complex that recognize and bind defined promotor sequences, thereby dictating cellular transcriptional programs (Merrick, 1993;Feklístov et al., 2014). Canonical sigma-70 family members engage promoters at -35 and -10 regions relative to the transcriptional start site, induce promoter open-complex formation, and initiate transcription (Zhang and Buck, 2015). In contrast, the structurally unrelated sigma-54 family members recognize unique -22 and -12 promoter regions; however, the formation of a repressive fork junction structure near -12 prevents the recruited holoenzyme from forming an open complex unassisted (Rappas et al., 2007;Zhang and Buck, 2015). Conversion to an open complex requires energy transfer from unique bacterial enhancer-binding proteins (EBPs, or sigma-54 activators) that bind conserved enhancer sequences upstream of the promoter. Formation of a hairpin-like structure, assisted by DNA-bending proteins such as integration host factor (IHF), allows enhancer-bound EBPs to interact with the promoter-bound sigma-54 closed complex (Zhang and Buck, 2015). Activator-mediated ATP hydrolysis results in loss of the fork junction structure and isomerization to an open complex, leading to transcription initiation.

Nie et al. used comparative genomics of 57 species from the *Clostridiales* order to reconstruct sigma-54 regulons and identify EBPs and their regulatory modules (Nie et al., 2019). Of the 24 predicted EBPs detected in *C. difficile*, the roles of two, CdsR and PrdR, involved in cysteine and proline catabolism, respectively, were experimentally validated (Bouillaut et al., 2013;Gu et al., 2017). To comprehensively characterize the SigL regulon, Soutourina et al used *in silico* analysis, transcriptome analyses and transcription start site (TSS) mapping of the laboratory-adapted *C. difficile* strain 630Δ*erm* (Soutourina et al., 2020). These studies revealed that the SigL regulon includes genes involved in the oxidation and reduction of amino acids to the corresponding organic acids (Stickland metabolism). Stickland reactions generate ATP and NAD^+^, and regulate *C. difficile* toxin production and virulence (Stickland, 1934;Neumann-Schaal et al., 2019).

To expand these observations to clinically-contemporary isolates, and further characterize the role of SigL in *C. difficile* biology, we generated isogenic insertion mutants (*sigL::erm*) in two genetically-divergent strains of human and veterinary origin: BI-1 (flagellated) and CDC1 (non-flagellated), belonging to the lineages (Ribotypes) RT027 and RT078, respectively. The epidemic-associated RT027 strains are often responsible for the most severe human *C. difficile* infections (CDI). RT078 strains, in contrast, while sharing several attributes with RT027 strains, are frequently associated with livestock infections (Elliott et al., 2017). We performed phenotypic assays and comparative proteomics studies on parental and *sigL::erm* derivatives. SigL modulated the *C. difficile* proteome in a temporal manner, and loss of *sigL* had profound, but contrasting and pleiotropic effects on the two strains. Our studies support a key role for SigL in *C. difficile* gene regulation, including strain-specific roles in the modulation of virulence factor expression, central metabolic functions, and sporulation.

## Results

### *C. difficile* encodes a unique sigma-54 ortholog

The *C. difficile* BI-1 (RT027) *sigL* ortholog is located immediately downstream from a putative purine biosynthesis locus (Fig. 1A). The encoded 454 amino acid protein includes conserved sigma-54 DNA-, RNA polymerase-, and transcriptional activator-binding domains. *sigL* is conserved among all sequenced *C. difficile* genomes, and the predicted amino acid sequence is identical between BI-1 and CDC1 strains.

**Figure 1.**
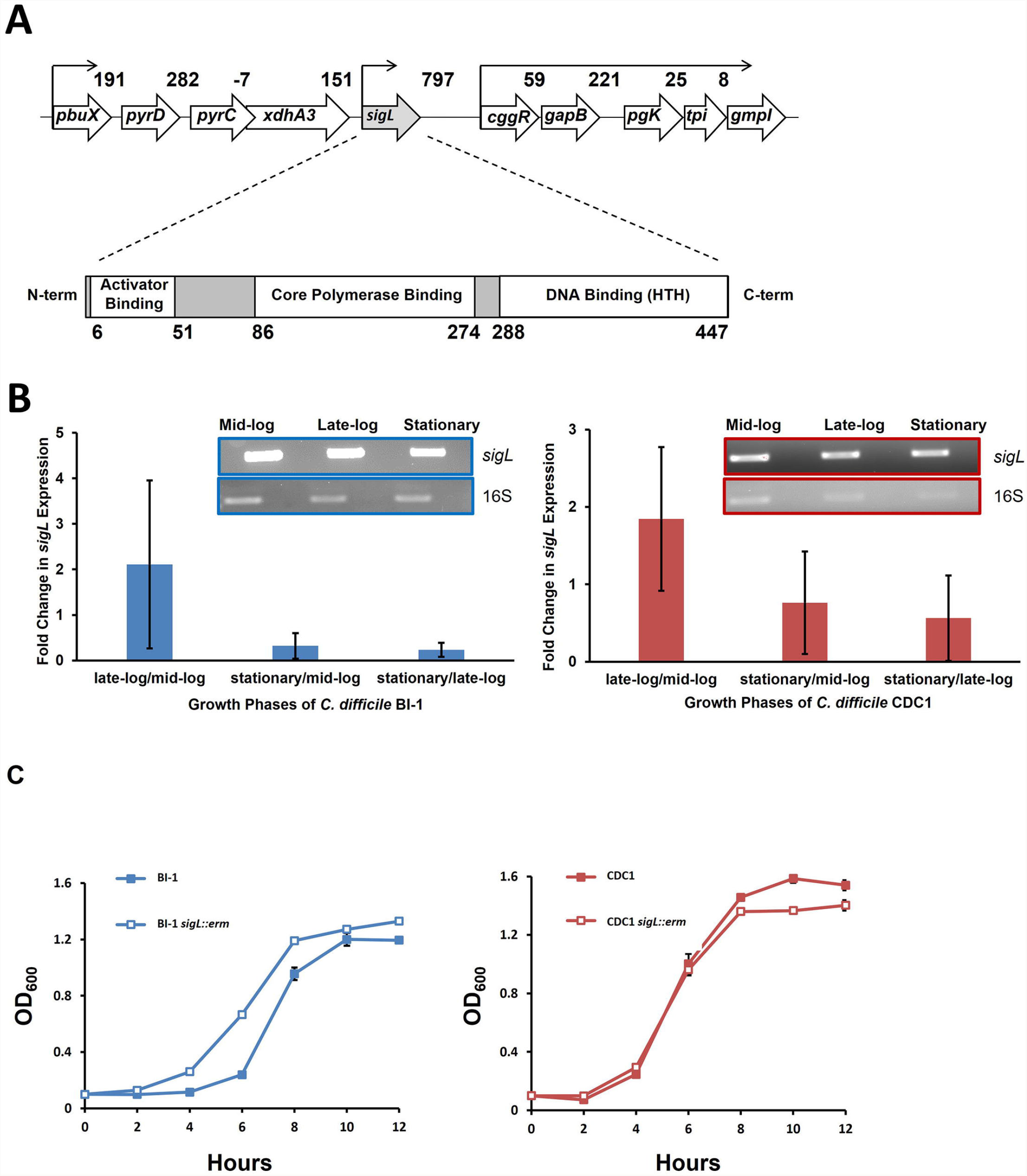
*sigL* loci, expression, and growth kinetics in *C. difficile* BI-1 and CDC1. **(A)**. *C. difficile sigL* locus, showing predicted promoters, based on strain BI-1 genome sequence, and SigL conserved binding domains. Numbers in the upper schematic represent intergenic distances, and numbers below the lower schematic indicate amino acids. **(B)**. Quantitative and semi-quantitative (inset) RT-PCR of *sigL* in BI-1 and CDC-1 across multiple growth phases. *16s* was used as a housekeeping control for qRT-PCR. Fold changes in expression represent the average of three biological replicates with standard deviation **(C)**. Growth of BI-1 and CDC1 strains compared to respective *sigL::erm* mutants. Cultures were grown in triplicate.

### *C. difficile sigL* is temporally expressed and is not essential for growth

During *C. difficile* growth, *sigL* expression was highest in late-exponential phase, and lowest in early growth- and stationary phases. These expression trends were similar in BI-1 and CDC1 strains (Fig. 1B). To interrogate the contribution of *sigL* to *C. difficile* biology, a lactococcal retrotransposition approach was used to generate isogenic mutants and corresponding complements in strains BI-1 and CDC1 (Kuehne and Minton, 2012) (Figure S1). While the *sigL::erm* strains did not exhibit viability defects, they displayed modest, distinct, and reproducible differences in growth kinetics relative to the isogenic parent strains (Fig. 1C).

### SigL regulates the abundances of diverse proteins

Soutourina and colleagues identified thirty SigL-dependent promoter sequences controlling 95 genes in the *C. difficile* strain 630 genome. Twenty-four of the identified genes or operons were adjacent to enhancer-binding protein (EBP) genes involved in sigma-54-dependent activation (Soutourina et al., 2020). They also noted that transcription levels of genes or operons in 12 of the 30 putative *sigL* promoter sequences were downregulated in microarray analysis of the *C. difficile* 630Δ*erm sigL::erm* mutant (GEO accession GPL10556). We used the 12 putative *sigL* promoter sequences whose genes or operons were downregulated to generate a *sigL* positional weight matrix (PWM); this was used for promoter-proximal SigL binding site determination by employing the Find Individual Motif Occurrences (FIMO) tool (Grant et al., 2011).

Consistent with Soutourina et al (2020), FIMO identified all 30 putative *sigL* promoter sites in both the 630 and BI-1 genomes, and notable differences from these two strains in the RT078 isolate, CDC1 (Table 1, Table S1). First, only 28 of the 30 sites were conserved in CDC1; CD0166-CD0165 (peptidase, amino acid transporter) and CD2699-CD2697 (membrane protein, peptidase) operons and their respective *sigL* promoter sequences were either missing or disrupted relative to 630 and BI-1 (Table 1). Second, the CD0284-CD0289 operon (PTS Mannose/fructose/sorbose family) exhibited a substantially lower FIMO score compared to 630 and BI-1, with an atypical G residue at position 18 of the PWM. Similarly, CD3094 (sigma-54-dependent transcriptional regulator) has an atypical G residue at position 17 in the CDC1 promoter site (Table 1). Third, all three strains exhibited unique insertions/deletions in the *sigL*-dependent *prd* operon, linked to proline metabolism (Figure S2). CD3246, a putative surface protein, is inserted between *prdC* and *prdR* in *C. difficile* 630, and is absent or truncated in BI-1 and CDC1. *prdB*, encoding a proline reductase, is present in 630 and BI-1 but truncated and missing a functional domain in CDC1 (Figure S2). Of the previously identified EBPs for *sigL*-dependent promoters, all 24 identified in *C. difficile* 630 (Nie et al., 2019;Soutourina et al., 2020) were conserved in BI-1, while two EBPs (CD0167 and CD2700) were missing in CDC1 (data not shown).

**Table 1.**
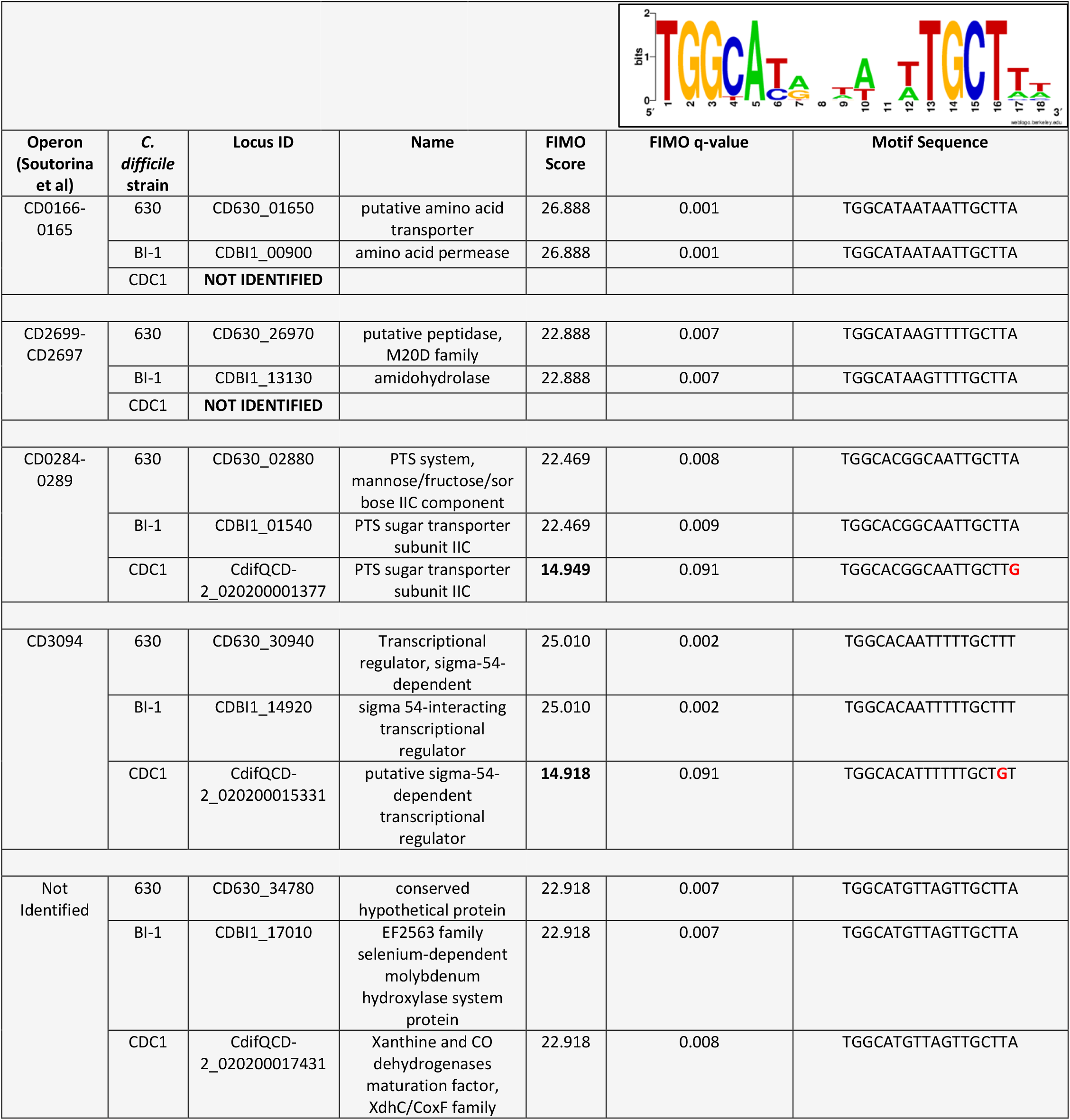
*sigL* positional weight matrix and *sigL* motif sequences with FIMO scores in *C. difficile* strains 630, BI-1 and CDC1. *sigL*-dependent operons identified by Soutorina et al are listed in the first column, with the exception of CD630_34780 identified in our study (bottom row). Associated strains, loci, names, FIMO scores/q-values and motif sequences are listed for the identified motifs. Two loci are not identified in CDC1 (column 3). Lower FIMO scores are bolded, and atypical nucleotides in the motif sequences are shown in red.

To gain a comprehensive appreciation of SigL-dependent regulation in ribotype RT027 and RT078 *C. difficile* strains, total protein abundances of BI-1, CDC1, and their respective *sigL::erm* derivatives were quantitated at mid-exponential (OD_600_ 0.6), late-exponential (OD_600_ 0.8), and stationary (OD_600_ 1.0) growth phases using quantitative mass spectrometry-based proteomics, and multiple stringent statistical tests. Overall, relative to the isogenic parent strains, more proteins were differentially abundant in CDC1 *sigL::erm* than in BI-1 *sigL::erm* across all three growth phases, but especially during late-exponential growth (Figure 2, Table S2). For BI-1, 335 of 1426 identified proteins were altered in abundance in *sigL::erm* in at least one growth phase relative to WT. This corresponds to 24.9% of all expressed proteins for this strain. For CDC1, 516 proteins were altered in abundance in *sigL::erm*, corresponding to 73.5% of all expressed proteins for this strain. There was also a marked downregulation of proteins implicated in isoleucine fermentation and short chain fatty acid metabolism in *sigL*::*erm* for both strains, and across all growth phases (Table S2). Based on the markedly different protein profiles between the two strains, we further explored strain-specific SigL-dependent regulation of *C. difficile* physiology, including cell surface-related phenotypes, sporulation kinetics, biofilm formation, and toxin production.

**Figure 2.**
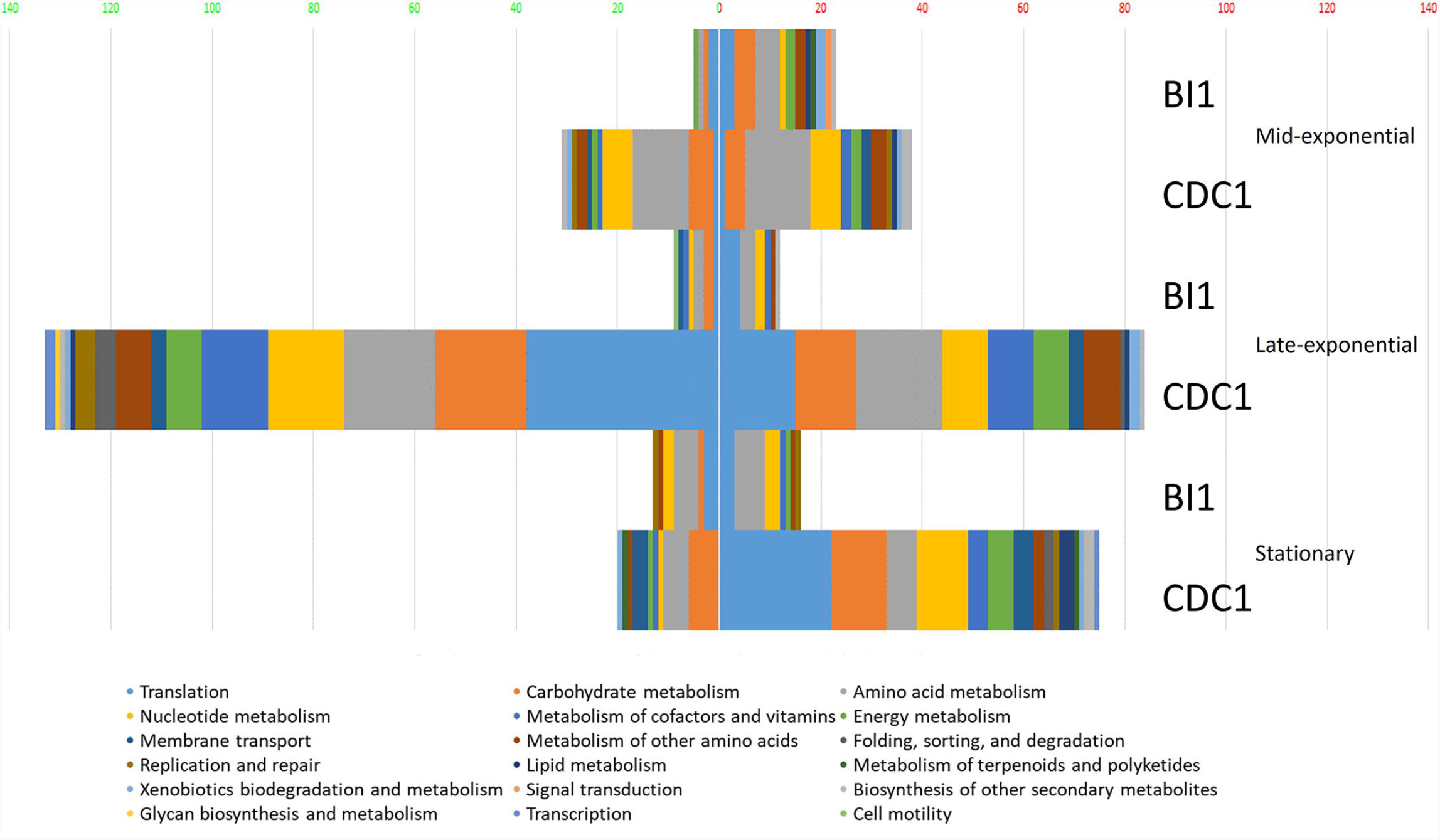
Functional classification of dysregulated proteins in *C. difficile* strains BI-1 and CDC1 resulting from *sigL* disruption. Functional classification analysis was performed on proteins identified as altered in *sigL* disruption mutants using an arbitrary fold change cut-off of 2. The x-axis is the number of dysregulated proteins. Numbers on the left in (green) are downregulated, on the right (red) are upregulated. The colors of the columns represent different functional categories based on RAST classification system.

### *sigL* mutants display an altered cell surface

Relative to the BI-1 WT strain, *sigL::erm* had 47 surface-associated proteins with altered abundance (21 upregulated, 28 downregulated), and for CDC1, *sigL::erm* had 61 surface-associated proteins with altered abundance (44 upregulated, 34 downregulated). Proteomics findings also revealed that dysregulated molecules included those predicted to be involved in S-layer display and host-cell adhesion. Therefore, we probed the surface properties of the *sigL::erm* mutants relative to the respective isogenic parent strains.

PSII, an antigenically-important surface polysaccharide unique to *C. difficile* strains, is hypothesized to assemble on the inner face of the cell membrane, and is translocated to the cell surface via the action of a flippase, MviN (Willing et al., 2015;Chu et al., 2016). Perturbations in PSII biogenesis and/or cell surface display result in diminished extractability of this molecule (Chu et al., 2016). Immunoblotting revealed diminished surface-extractable PSII in BI-1 *sigL::erm* compared to the isogenic parent strain. In contrast, however, PSII was more readily released from CDC1 *sigL::erm* strain relative to the isogenic parent (Figure 3). Thus, SigL influences expression, export and/or assembly of key *C. difficile* surface molecules in a strain-specific manner.

**Figure 3.**
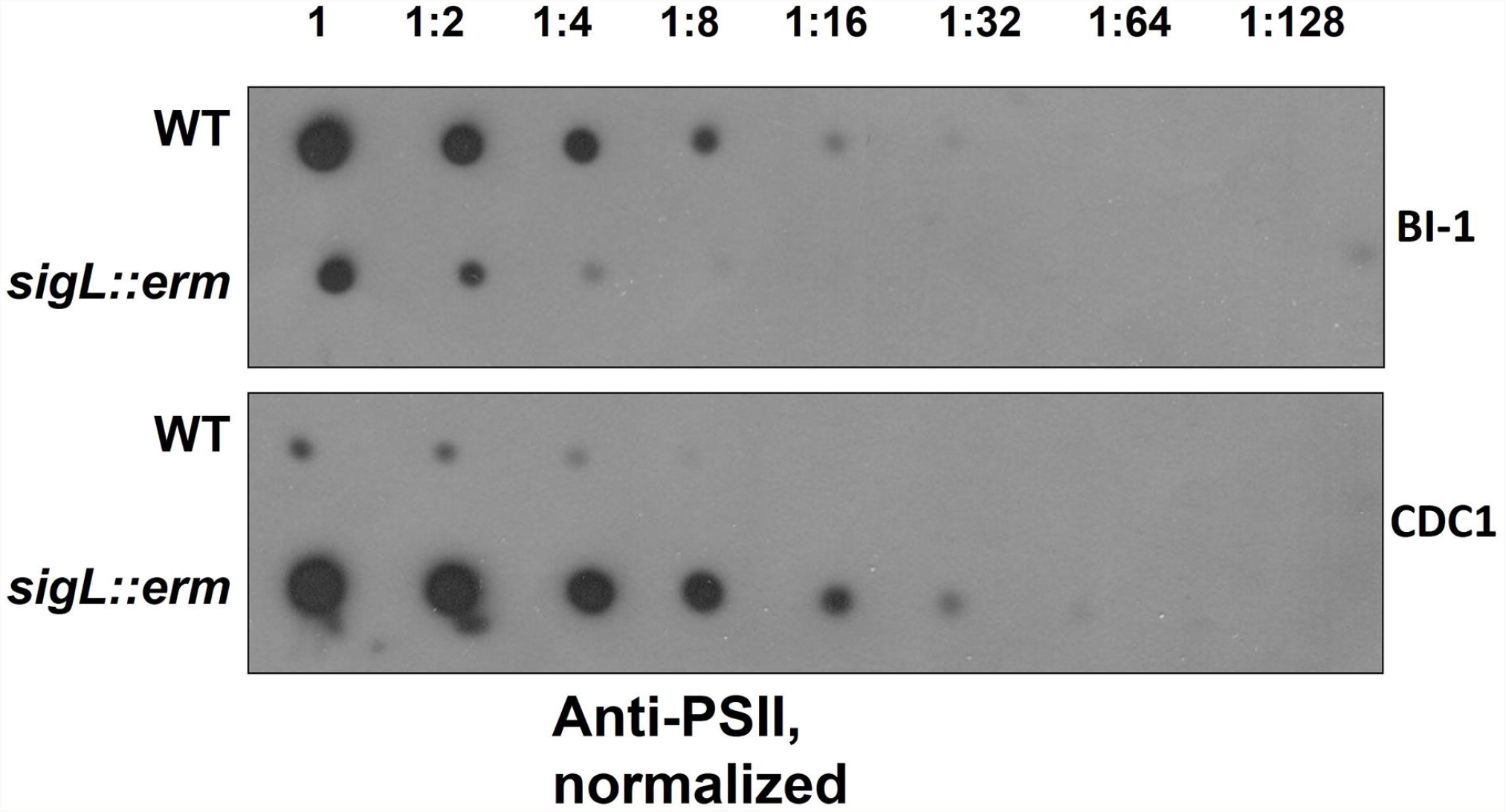
PSII abundance in *sigL::erm* differs with respect to wild-type strain BI-1 and CDC1. Serially-diluted cell extracts of WT and *sigL::erm* derivatives (normalized to OD600=1.2 in BHI) were spotted on PVDF membranes and immunoblotted using PSII-specific antiserum.

Changes in bacterial surface architecture impact autolysis susceptibility; this has implications for toxin release in *C. difficile* (Tsuchido et al., 1990;Wydau-Dematteis et al., 2018;Mayer et al., 2019). Surface composition dictates the ease with which detergents, such as Triton X-100, interact with lipoteichoic acid from Gram-positive bacterial cell walls, which in turn influences activation of autolytic enzymes (Komatsuzawa et al., 1994;Fujimoto and Bayles, 1998). To establish the influence of SigL on *C. difficile* cell surface properties, we quantitatively measured rates of autolysis between wild type and *sigL::erm* mutants of both BI-1 and CDC1. While autolysis rates were similar between the BI-1 WT and *sigL::erm* strains at early time points, the mutant demonstrated decreased autolysis at later time points. In contrast, the CDC1 *sigL::erm* strain exhibited a more rapid rate of autolysis than the WT strain, and underwent complete autolysis by 90 minutes (Figure 4A, B).

**Figure 4.**
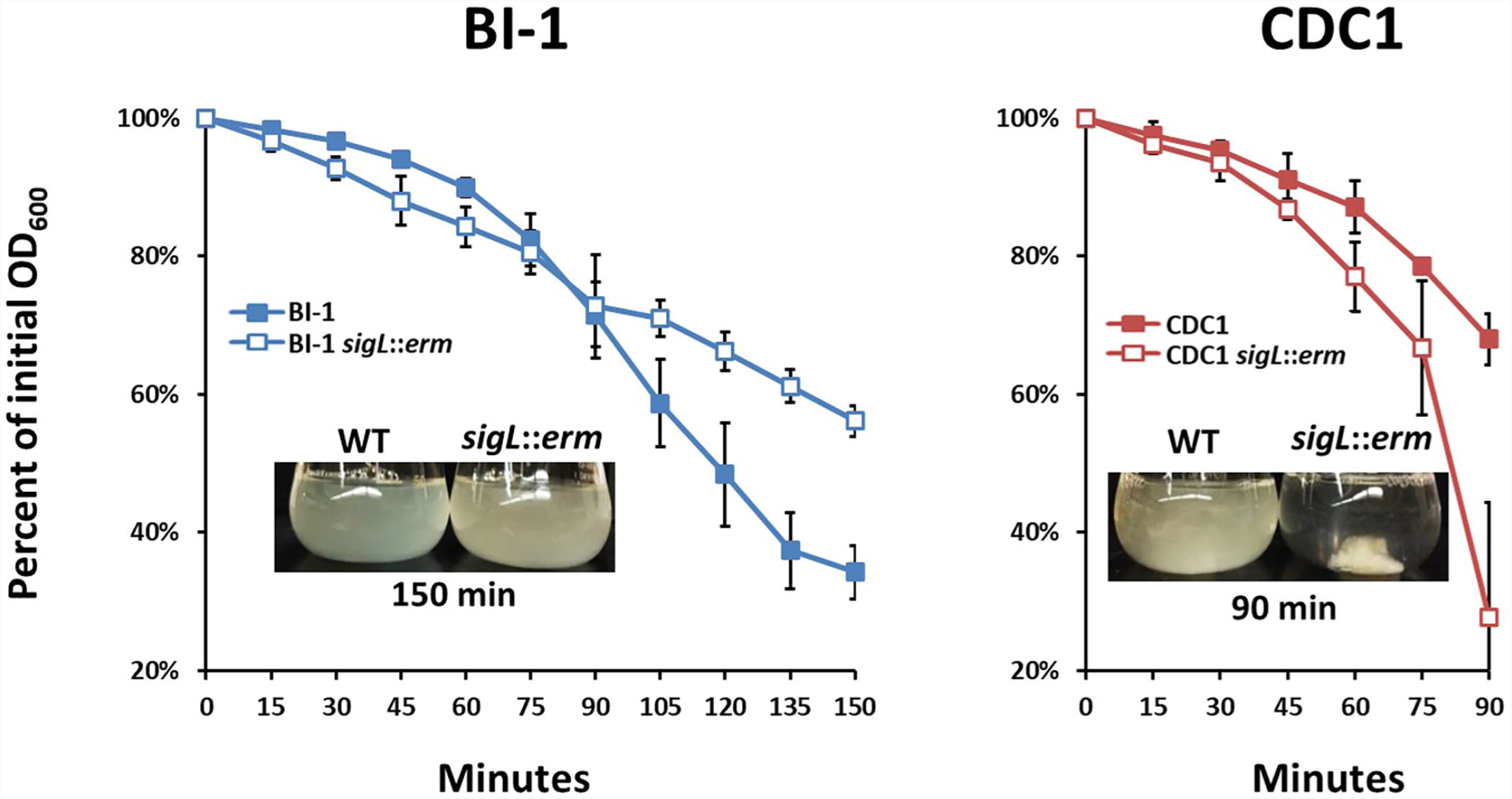
Autolysis rates of *C. difficile* strains and *sigL* mutants differ between BI-1 and CDC1. Cultures were grown to OD_600_ level of approximately 1.2, and OD_600_ values were measured as percent of initial OD over time following the addition of Triton X-100. Visualization of autolysis in BI-1 and CDC1 compared to respective *sigL::erm* mutants are show as insets. Each assay was performed in biological triplicate.

### Loss of SigL perturbs *C. difficile* aggregation in a strain-specific manner

Changes in cell surface protein composition, including variation in charge and hydrophobicity can dictate bacterial aggregation which, in turn, can influence virulence (Misawa and Blaser, 2000). Therefore, we monitored aggregation rates of both BI-1 and CDC1 *sigL::erm* mutants during growth in BHI. While the aggregation rate of *C. difficile* BI-1 and the isogenic *sigL::erm* remained unchanged throughout growth (Figure 5A), CDC1 *sigL::erm* was more aggregative than the parent WT strain during late-exponential and stationary growth phases (Figure 5A). These phenotypes were independently verified via autoagglutination assays using plate-grown cultures. Again, CDC1 *sigL::erm* agglutinated significantly more than the cognate wildtype strain at multiple time points (inset, Figure 5B). Autoagglutination rates of the BI-1 *sigL::erm* were not different from wild type with the exception of two time points at 8 and 10 hours (inset, Figure 5B).

**Figure 5.**
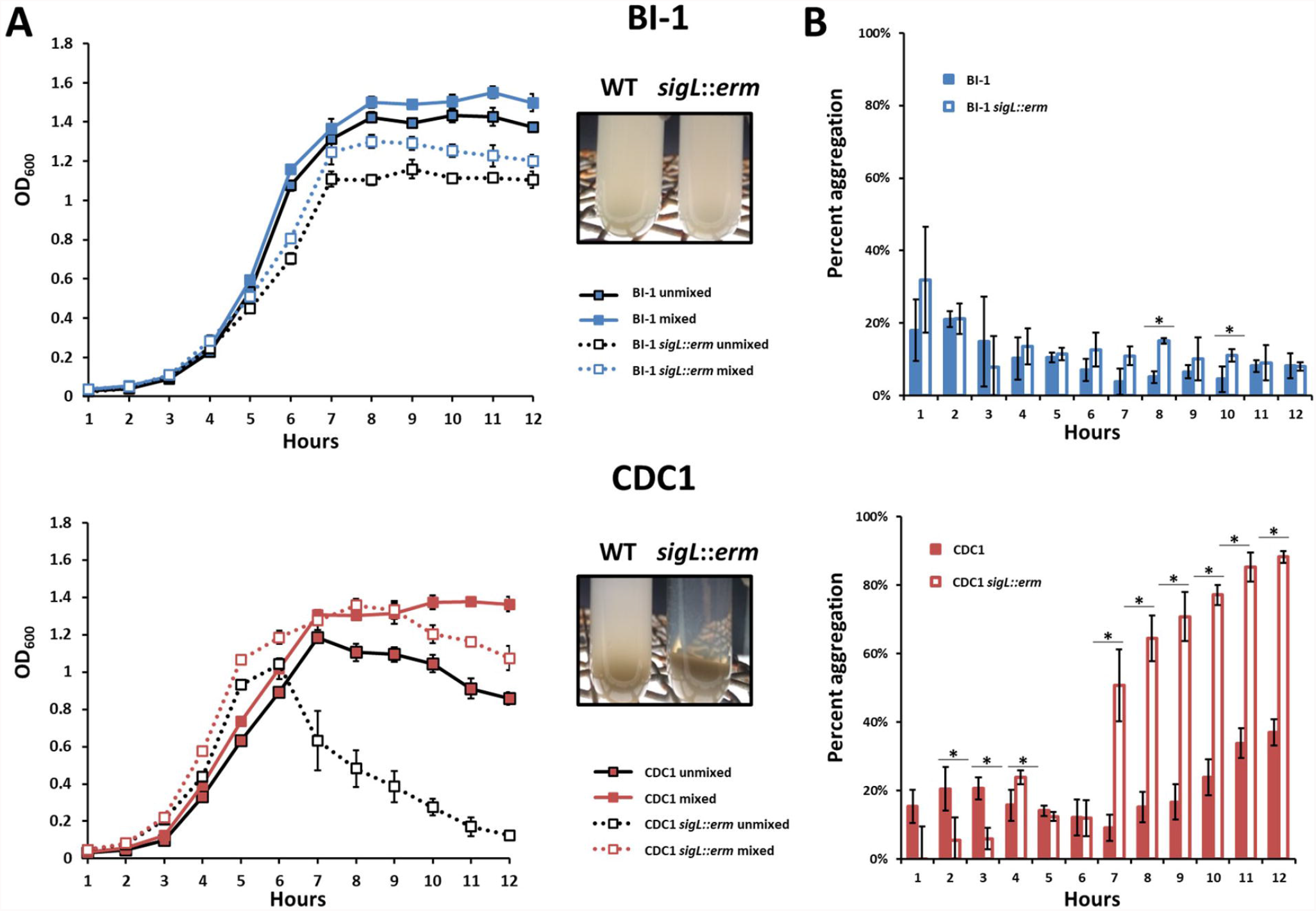
Aggregation of *C. difficile* strains and respective *sigL::erm* mutants differs between BI-1 and CDC1. **(A)**. OD_600_ values of top 1 mL of culture (unmixed) and a replicate vortexed culture sample (mixed) were read hourly to determine sedimentation curves. (Inset). Visualization of aggregation in BI-1 (top) and CDC1 (bottom) WT and *sigL::erm* strains. and percent aggregation as a calculation of (1-OD600 unmixed)/(OD600 mixed) (Right) of BI-1 and BI-1 *sigL::erm*. **(B)**. Percent aggregation of *C. difficile* strains and respective *sigL::erm* mutants, calculated as the difference in OD_600_ values between whole (mixed) culture and top 1 mL (unmixed) culture, as a percentage of the mixed culture OD_600_ value. Cultures were grown in biological triplicate. WT and *sigL::erm* pairs were compared at each time point using Student’s t-test. *p<.05

### *C. difficile sigL* mutants exhibit altered propensity to form biofilms

Altered surface properties can influence the ability to form biofilms; in *C. difficile*, this can impact pathogen persistence in the gut (Pantaléon et al., 2015;Fernández Ramírez et al., 2018;Chamarande et al., 2021;Janež et al., 2021;Kumari and Kaur, 2021). Disruption of SigL had no discernable impact on the ability of BI-1 strains to form biofilms (Figure 6A). In contrast, CDC1 *sigL::erm* had decreased biofilm density relative to the isogenic parent strain (Figure 6B). Complementation restored the CDC1 phenotype (Figure S3A), and overexpression of SigL enhanced the ability of the CDC1 strain, but not BI1, to form biofilm (Figure S3B,C).

**Figure 6.**
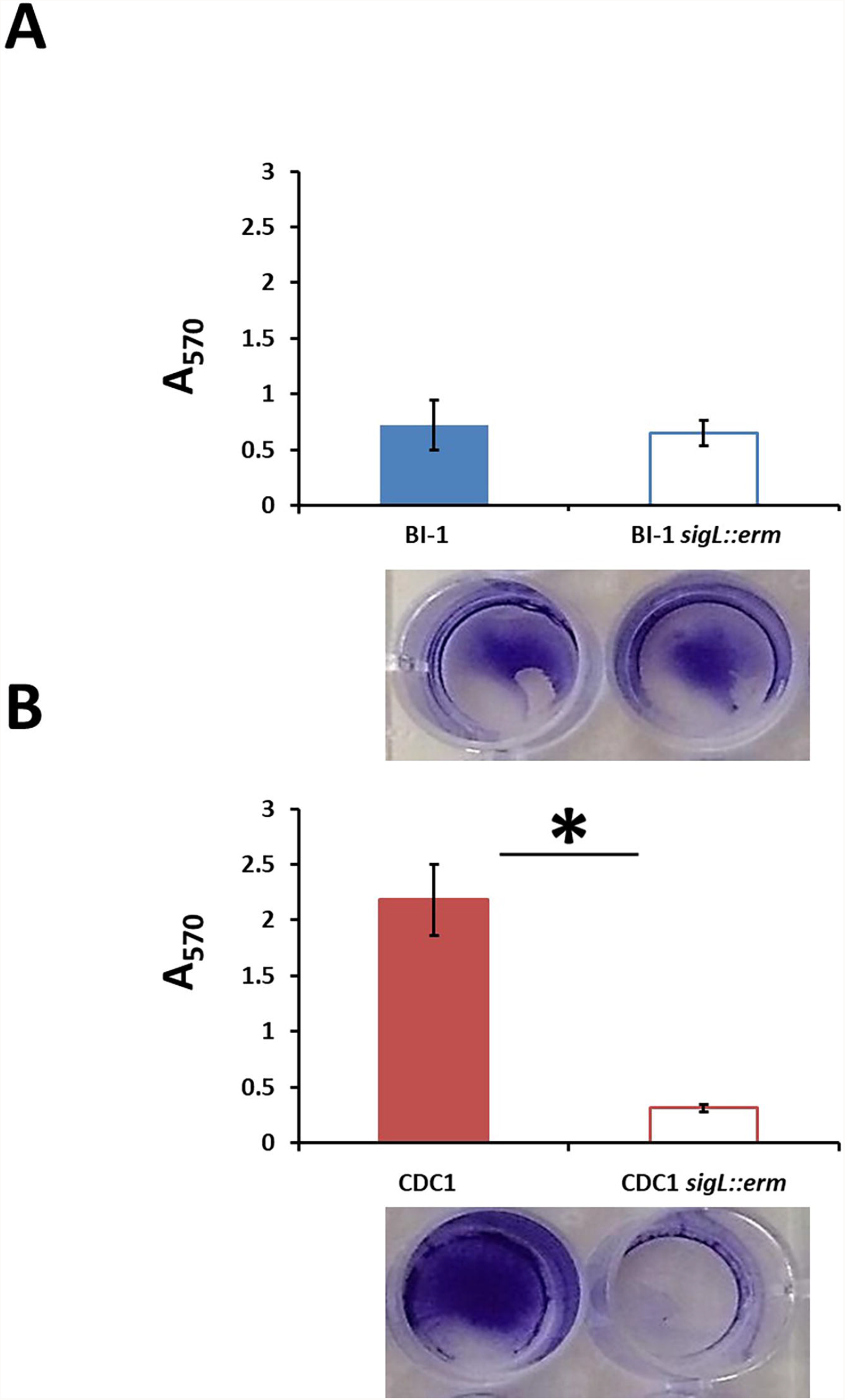
Biofilm formation is reduced in CDC1 *sigL::erm*, but not in BI-1 *sigL::erm*. Biofilm was quantified by the release of crystal violet retained by the *C. difficile* biomass after 72 hours. Absorbance was measured at 570 nm. **(A)**. Biofilm formation in BI-1 and BI-1 *sigL::erm*. **(B)**. Biofilm formation in CDC1 and CDC1 *sigL::erm*. Samples were grown in biological triplicate and compared using Student’s t-test. *p<.05

### *sigL* mutants exhibit altered levels of attachment to cultured human colonic intestinal epithelial cells

Variations in cell surface composition leading to alterations in phenotypic properties including aggregation can affect bacterial attachment to host cells (Burdman et al., 2000;Merrigan et al., 2013;Foster, 2019), and this has profound implications for *C. difficile* host establishment (Vedantam et al., 2018). We therefore compared the adherence of parental and *sigL* mutant strains to Caco-2_BBe_ intestinal epithelial cells. Two different growth phases were chosen for analysis representing differential protein expression in *sigL::erm* strains (Figure 7): OD_600_=0.4, mid-exponential, and OD_600_=1.2, stationary. For mid-exponential phase bacteria, compared to the respective parent strains, BI-1 *sigL::erm* exhibited diminished levels of attachment, whereas CDC1 *sigL::erm* attachment was more robust (Figure 7A). For bacteria at stationary phase, there was no apparent difference in attachment between the WT and the mutant in the BI-1 background, but CDC1 *sigL::erm* was less adherent to host cells relative to the isogenic WT strain (Figure 7B). Thus, the contribution of SigL toward the ability of *C. difficile* to attach to biotic surfaces is both strain- and growth phase-dependent.

**Figure 7.**
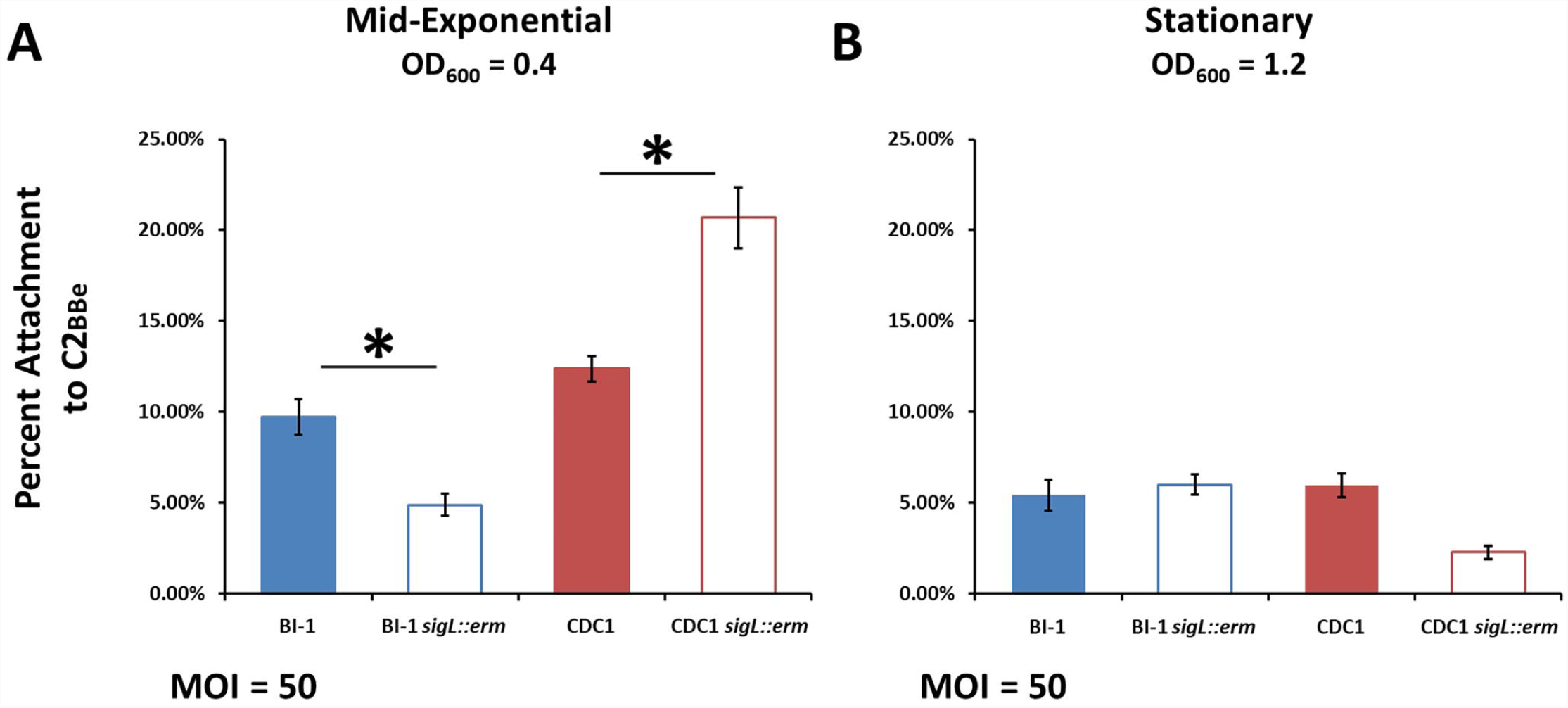
Host cell attachment in *sigL::erm* is phase- and ribotype-dependent. *C. difficile* cultures were grown to mid-exponential or stationary phase and added to Caco-2_BBe_ intestinal epithelial cell monolayers. Percent attachment was calculated by adhered CFUs as a percentage of the initial inoculum CFUs. **(A)**. Attachment to Caco-2_BBe_ intestinal epithelial cells in mid-exponential phase. **(B)**. Attachment to Caco-2_BBe_ intestinal epithelial cells in stationary phase. The assays were conducted in biological triplicate and samples were compared using ANOVA and Tukey’s HSD post-hoc test. *p<.05

### Loss of *sigL* alters sporulation kinetics in an RT078 background

*C. difficile* sporulates in response to a combination of unfavorable extracellular environment and nutritional/metabolic deprivation, and sporulation efficiency can impact bacterial gut persistence (Paredes-Sabja et al., 2014;Shen et al., 2019). We next assessed if loss of SigL influences *C. difficile* sporulation kinetics. BI-1 and BI-1-*sigL::erm* formed spores at comparable levels over a period of five days, indicating that SigL does not influence sporulation processes in this strain (Figure 8A). In contrast, CDC1 *sigL::erm* formed significantly greater number of spores than the WT strain at 24 and 48 hours, but the difference resolved at later time points (72-120H) (Figure 8B).

**Figure 8.**
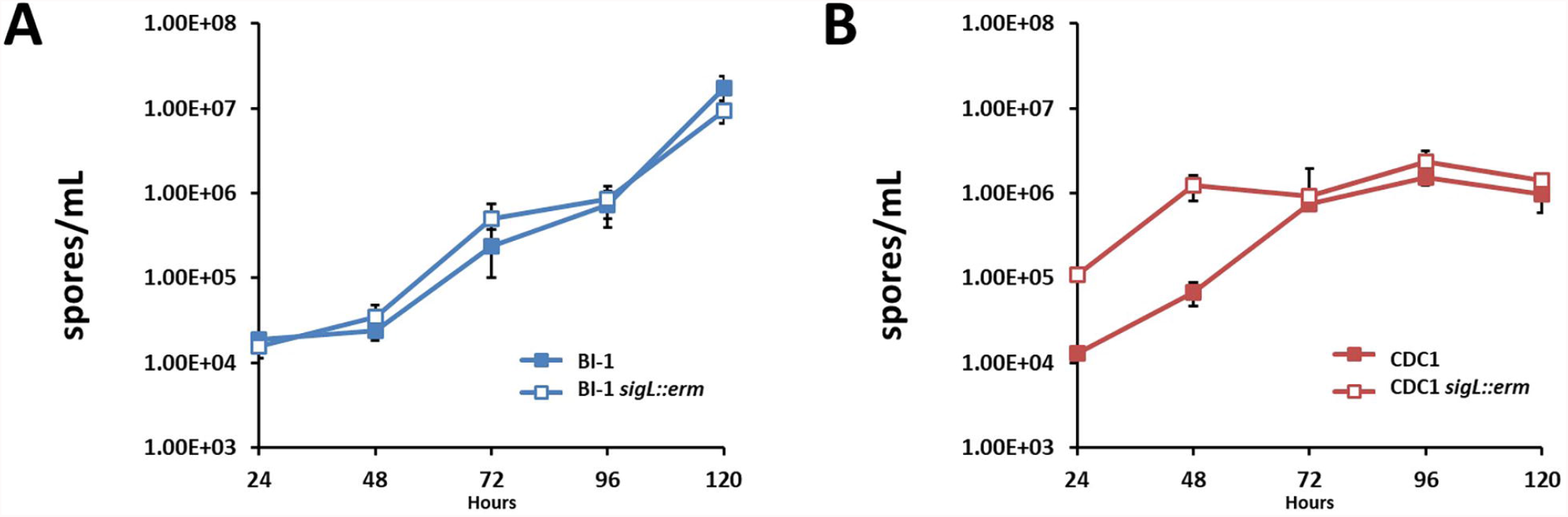
Sporulation kinetics are altered in *sigL::erm* in a ribotype-dependent manner. Spores were enumerated by heat shocking and plating cultures on BHIS-Taurocholate at each time point. **(A)**. Spore enumeration over 120 hours in BI-1 and BI-1 *sigL::erm* strains **(B)**. Spore enumeration over 120 hours in CDC1 and CDC1 *sigL::erm* strains. Cultures were grown in biological triplicate.

We next assessed if the increased sporulation phenotype of CDC1 *sigL::erm* could be reversed by plasmid-complementation. Plasmid (vector)-harboring WT- and mutant strains recapitulated the sporulation phenotypes for CDC1 though differences in overall kinetics were observed (likely due to the presence of the antibiotic used for plasmid maintenance). This was reversed by a *sigL-*encoding plasmid (Figure S4). Spore production was also reduced in BI-1 with plasmid complementation. Plasmid-driven SigL overexpression depressed sporulation of both the CDC1- and BI-1-*sigL::erm* mutants, as well as the corresponding WT strains (Figure S5).

### *C. difficile* BI-1 motility is not altered by *sigL* inactivation

Sigma-54 is a widely recognized as an important regulator of flagellar biosynthesis in many Gram-negative species, but its role is non-uniform among different families of organisms (Tsang and Hoover, 2014). To evaluate the role of SigL in regulation of flagellar synthesis in *C. difficile*, motility assays were performed using the motile strain BI-1. CDC1 is aflagellate due to the loss of a substantial part of the flagellar locus encoding early structural flagellar genes (Stabler et al., 2009), a hallmark of the RT078 ribotype, and was used as a negative control. In contrast to other bacterial systems whereby motility is influenced by sigma-54 expression, the motility and flagellar expression of *C. difficile* BI-1 was not significantly affected by *sigL* inactivation (Figure S6).

### *C. difficile* secreted toxin levels are altered in *sigL* mutants

*C. difficile* toxin expression is influenced by both metabolic factors and genetic determinants (Bouillaut et al., 2015;Martin-Verstraete et al., 2016;Ransom et al., 2018). Given the wide-ranging impact of SigL presence on *C. difficile* metabolic enzymes, we used immunoblot analysis to monitor TcdA and TcdB in WT and mutant culture supernatants. Loss of SigL dramatically increased secreted TcdA and TcdB levels in BI-1, but decreased the levels of the two toxins in the CDC1 background (Figure 9). This was confirmed by toxin ELISA (Figure S7). Correspondingly, complementation decreased toxin expression by BI-1 *sigL::erm*, but increased TcdA and TcdB levels in the CDC1 background (Figure S7).

**Figure 9.**
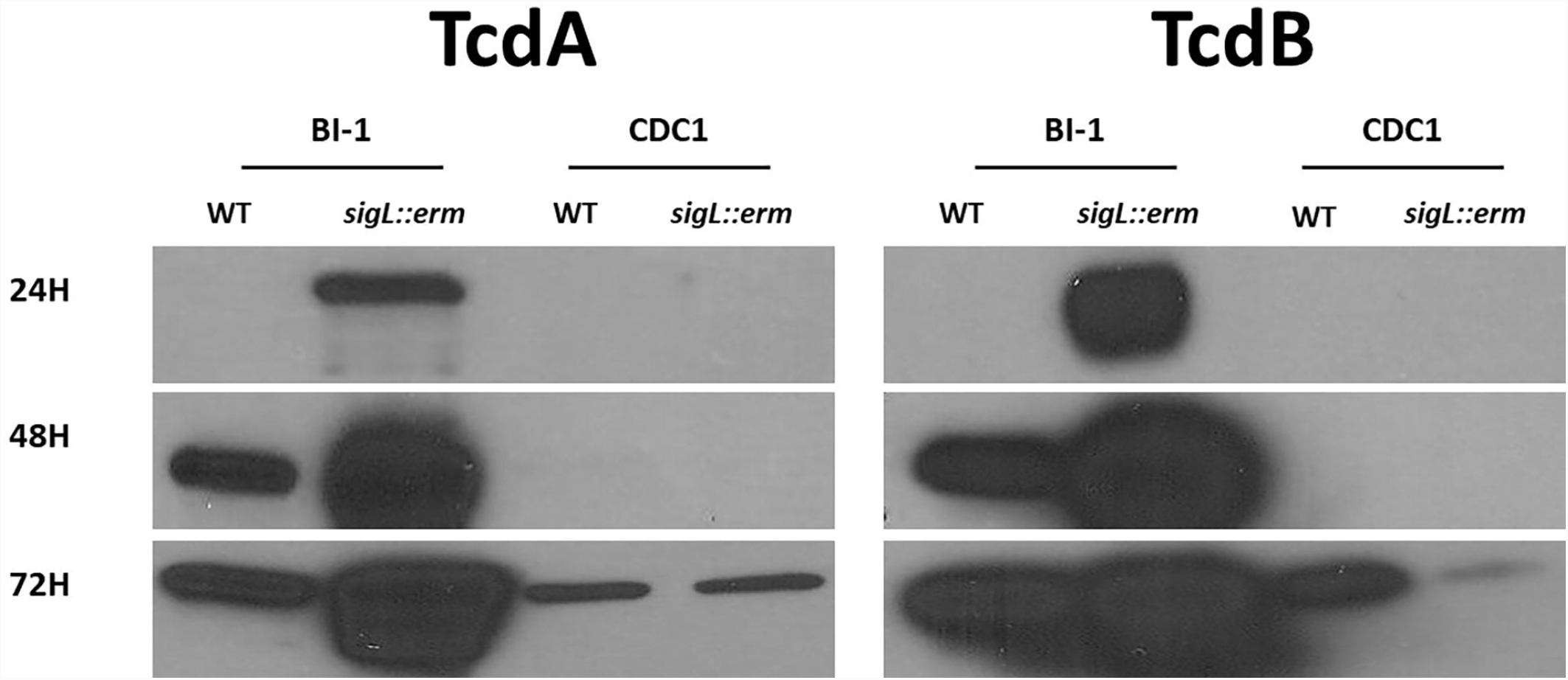
Toxin production is differentially altered in *sigL::erm* in BI-1 and CDC1 backgrounds. BI-1 and CDC1 *C. difficile* strains and respective *sigL::erm* mutants were grown for 24, 48, and 72 hours. Vegetative cells were removed by centrifugation and supernatant was assayed for toxin by immunoblotting using monoclonal antibodies against TcdA and TcdB. Blots are representative of three experiments.

## Discussion

Although rarely an essential protein, no unified functional theme has been identified concerning the sigma-54 dependent metabolic repertoire. This is in contrast with the sigma-70 counterpart, which often serves dedicated functions. Sigma-54-dependent proteins have pleotropic functions in bacterial systems, with roles ranging from carbohydrate and nitrogen metabolism to modification of cell surface structures including flagella and pili (Alm et al., 1993;Francke et al., 2011;Hayrapetyan et al., 2015). In Gram-positive organisms, the Sigma-54, SigL, regulates adaptation to environmental conditions including cold shock (Wiegeshoff et al., 2006), osmotic stress (Okada et al., 2006) and oxidative stress (Stevens et al., 2010), and is involved in amino acid catabolism (Debarbouille et al., 1999), acetoin cycle regulation (Ali et al., 2001;Francke et al., 2011), and flagellar synthesis (Hayrapetyan et al., 2015).

A previous report revealed that *C. difficile sigL* transcript levels were highest during the stationary phase of growth (Karlsson et al., 2008); in this study, we observed maximal SigL expression during late-exponential phase. Due to its requirement for activation prior to transcription, it is likely *C. difficile* SigL function is regulated through the coordinated expression of EBPs, driving SigL transcription of target operons when physiologically necessary. The number of EBPs roughly correlates with genome size, with *C. difficile* strains 630 and BI-1 harboring 23 such genes, two of which are absent in CDC1 (Nie et al., 2019).

Promoters recognized by members of the sigma-54 family have highly conserved -22 and -12 regions upstream of the transcriptional start site, including a GC dinucleotide at position -12 and GG at position -24 (Barrios et al., 1999;Zhang and Buck, 2015). Barrios et al identified 85 SigL promoter sequences in multiple bacterial species, with the consensus sequence ‘TTGGCATNNNNNTTGCT’ used to inform more recent studies to search for promoter candidates in *Clostridiales* genomes (Barrios et al., 1999;Nie et al., 2019). While *C. difficile* SigL interaction with specific promoters was described previously (Nie et al., 2019;Soutourina et al., 2020), the relative strength of the promoters, or the essentiality of specific residues in the promoter for interaction, has not been defined.

We identified SigL-dependent promoter sequences in BI-1 and CDC1 strains in agreement with previous studies in *C. difficile* 630, with SigL-dependent regulation of genes involved in amino acid metabolism and sugar transport, in addition to novel promoter sequences not identified in previous 630 studies (Soutourina et al., 2020). In BI-1, a promoter sequence for an EF2563 family selenium-dependent molybdenum hydroxylase (CD630_3478/CDBI1_17010) was uniquely identified. This same EF2563 protein was identified in CDC-1 (CdifQCD-2_020200017431) (Table 1, Table S1). Other strain-specific differences noted in the repertoire of SigL regulated genes included the absence of two SigL-dependent promoters in CDC1 compared to BI-1 and 630, corresponding to an amino acid transporter and a membrane protein/peptidase respectively. Additionally, a G residue replaces the typical A or T at position 18 in the positional weight matrix (PWN) for the CD0284-CD0289 operon (PTS Mannose/fructose/sorbose family) and position 17 for the CD3094 operon, resulting in a lower FIMO score suggestive of decreased SigL interaction. These unique promoter features noted for BI-1 and CDC-1 extended to other published sequences of strains belonging to the RT027 and RT078 lineages, respectively.

In our proteomics analysis of BI-1 and CDC1, we observed profoundly pleotropic, strain-specific, and growth-phase-specific SigL-dependent regulation. In late-exponential phase, 224 proteins were downregulated in the CDC1*::erm* strain, compared to only 76 downregulated proteins in the BI-1 background for the same growth phase (Table S2). Our proteomics data showed that several hydroxyisocaproyl-CoA dehydratase (Had) proteins, associated with L-leucine fermentation, were downregulated in *sigL::erm* in both strain backgrounds and across all growth phases, consistent with established roles for sigma-54 in amino acid fermentation (Gardan et al., 1995;1997;Debarbouille et al., 1999). Our data also show that SigL-dependent protein regulation is not only highly pleotropic within a strain, but widely differential between *C. difficile* 027 and 078 strains.

The SigL regulon is remarkably distinct between BI-1 (RT027) and CDC1 (RT078), and this is manifested via distinct phenotypes. Such differences have been shown for *C. difficile* strains R20291 (RT027) and 630Δ*erm* (RT012) specifically in the context of flagellar protein mutants (Baban et al., 2013), but comparative studies for sigma-54 mutants are lacking. A recent large-scale study identified evolutionary relationships between sigma-54 genes and the synthesis and transport of elements of the bacterial cell surface and exterior including exopolysaccharides, lipids, lipoproteins and peptidoglycan (Francke et al., 2011). In agreement, several dysregulated protein targets observed between the *C. difficile sigL* mutants were either components of, or associated with, the cell surface indicating dynamic compositional regulation by SigL. Phenotypic analysis of the *sigL* mutants of the two ribotypes confirms strain-specific surface alterations. Differential cell surface phenotypes with respect to aggregation, attachment to biotic surfaces, exopolysaccharide levels, and autolysis were observed.

Metabolic events and environmental conditions encountered during stationary phase determine whether *C. difficile* vegetative cells will form biofilms, produce toxin and/or initiate sporulation. In this work, strong correlation was observed between the expression of SigL, sporulation, biofilm, and toxin phenotypes. In CDC1, there is an inverse association between sporulation and toxin production: in CDC1 *sigL::erm*, sporulation levels are increased, and biofilm and toxin production are decreased relative to the parent strain. In BI-1, the relationship appears more complex. Sporulation and biofilm formation are not influenced by the loss of *sigL* expression in the BI-1 background; therefore, the reason for the accompanying increase in toxin production in the BI-1 *sigL* mutant remains unknown, but our observed strain-specific differences in SigL-dependent operons may yield some insight. The *prd* locus is associated with proline metabolism, which along with other amino acids can be metabolized by *C. difficile* via Stickland reactions. This amino acid metabolism is associated with reduced toxin production (Bouillaut et al., 2013). We identified a SigL-dependent promoter for the *prd* locus; therefore, a contributing factor to the increased toxin in BI-1 *sigL::erm* may be differences in SigL-dependent proline metabolism. Protein levels in members of the *prd* locus, including the proline reductase PrdA and proline racemase PrdF, were shown to be similarly decreased in *sigL::erm* in both the BI-1 and CDC1 backgrounds (Table S2), but interestingly, we observed a difference in the architecture of the *prd* locus in CDC1 compared to BI-1, with a truncation in *prdB* (Figure S2). How this truncation may impact proline metabolism and toxin production in CDC1 requires further investigation. There are also potential epistatic effects of other regulators, such as SigD, that may affect toxin production.

A recent study in *C. difficile* 630Δ*erm* showed major proteomic differences in two models of biofilm-like growth: aggregate biofilms and colony biofilms, and demonstrated that SigL was significantly induced in aggregate biofilms (Brauer et al., 2021). The proteomic and phenotypic differences in our respective 027 and 078 *sigL::erm* mutants may be partially explained by strain-specific differences in ability to aggregate and form biofilms, resulting in differential SigL induction. Indeed, WT BI-1 shows reduced baseline biofilm formation compared to WT CDC1 (Figure 6).

It is well-appreciated that strain-level variations may influence *C. difficile* virulence and epidemiology in fundamental ways, but less is known about the regulatory networks underlying these differences. Our studies suggest that the conserved regulator SigL has profoundly differential impacts on gene expression, including toxin production, in the predominantly human- and veterinary-associated *C. difficile* lineages RT027 and RT078, respectively. Future studies, including animal experiments, will be instructive in determining the consequences of these regulatory differences to disease establishment, progression, asymptomatic carriage, recurrence and/or response to treatment.

## Methods

### Bacterial Strains and Growth Conditions

All *C. difficile* strains utilized in this study (Table S3) were grown at 37°C using a Type B Coy Laboratory Anaerobic Chamber under anaerobic conditions in an 85% N_2_, 10% H_2_, 5% CO_2_ gas mixture. Bacteria were recovered from freezer storage in glycerol and routinely maintained on BHIS agar plates (Brain-Heart Infusion base media supplemented with 5g/L Yeast Extract and 0.1% L-Cysteine). All assays requiring liquid culture were performed using Bacto™ BHI unless otherwise indicated.

Antibiotics were used as necessary for *C. difficile*: (15μg/mL thiamphenicol, 20μg/mL lincomycin, 0.8μg/mL cefoxitin, 25μg/mL cycloserine) added to either BHI or BHIS. *E. coli* Top10 (Invitrogen) used for cloning and plasmid construction, and CA434 used for conjugation were routinely grown at 37°C on LB media using antibiotics as necessary: (20μg/mL chloramphenicol, 100μg/mL spectinomycin, 150μg/mL ampicillin). The *C. difficile* clinical isolates used in this study were from the Hines VA Hospital culture collection of Dr. Dale Gerding, or from the collection of Dr. Glenn Songer.

### *In silico sigL* sequence, promotor, and functional domain analysis

*C. difficile* genome sequences were maintained and annotated using Rapid Annotation using Subsystem Technology (RAST) bioinformatics web tools (Overbeek et al., 2014). *sigL* promotor prediction reported here was undertaken using Microbes Online web tools (Dehal et al., 2010). Functional domains of *C. difficile* SigL reported here were analyzed using Similar Modular Architecture Research Tool (SMART) (Letunic et al., 2015).

### Quantitative real-time PCR

*C. difficile* BI-1 and CDC1 strains were grown to mid-exponential (OD600 ∼0.6), late-exponential (OD600 ∼0.8), and stationary phase (OD600 ∼1.0). At each phase of growth, cultures were diluted 1:2 in RNA*later* RNA stabilization reagent (Sigma Aldrich) and incubated anaerobically for 1 hour, before pelleting and freezing bacteria at -80°C. RNA was extracted from cell pellets using the RNeasy Plus Mini Kit (Qiagen) according to manufacturer instructions, with modifications. Briefly, beat-beading tubes were filled 1/3 with beads and soaked overnight in 500 μl buffer RLT (Qiagen). Tubes were spun at full speed and excess buffer removed. Bacterial pellets were lysed by bead beating for 2 minutes in 600 μl buffer RLT and 2-Mercaptoethanol (10 μl/mL) per sample. Tubes were centrifuged at max speed for 3 minutes, and the supernatant was removed for RNA extraction according to manufacturer instructions. RNA samples were treated with DNase I to remove genomic DNA contamination, and RNA yields were quantified on a Nanodrop instrument. cDNA synthesis from the RNA samples was performed using the SuperScript III First Strand Synthesis System (Invitrogen) according to manufacturer instructions and yields were quantified using Nanodrop. Standard PCR was performed to confirm expression of *sigL* and *16s* (housekeeping gene), with Taq 2x Master Mix, using AC106 *sigL*-F/AC106 *sigL*-R and *16s*RT_F/*16s*RT_R primer pairs respectively (Table S4). Quantitative real-time PCR was performed using iTaq Universal SYBR Green Supermix (BioRad) on the StepOnePlus instrument (Applied Biosystems) according to manufacturer instructions. Expression fold changes were determined using ΔΔCT calculations.

### Generation and confirmation of *sigL* mutants

Isogenic *sigL* mutants were created in *C. difficile* BI-1 and CDC1 using the ClosTron system (Kuehne and Minton, 2012). Briefly, retargeted ClosTron introns were designed using the Perutka algorithm at Clostron.com and synthesized by DNA2.0 (DNA2.0, Melo Park, CA). The resulting constructs were electrotransformed into *E. coli* CA434 donor strain which was recovered on LB supplemented with spectinomycin and chloramphenicol. Isolated colonies were inoculated into BHIS broth supplemented with chloramphenicol and grown overnight at 37°C with shaking. One milliliter of culture was pelleted at 1500xg for 1 minute, washed in BHIS, centrifuged again for 1 minute at 1500xg, and transferred to an anaerobic chamber. The pellet was subsequently resuspended with 200μL of *C. difficile* overnight recipient culture and incubated overnight on BHIS agar. Growth was recovered in one milliliter of PBS, and 200μL aliquots were inoculated to BHIS(TCC) and incubated overnight at 37°C. Thiamphenicol-resistant colonies were isolated on BHIS supplemented with lincomycin to isolate potential integrant clones. Integration of the intron was confirmed by PCR using primers which detect both the integrated form of the intron, as well as primers AC55 *sigL*-F and AC56 *sigL*-R which amplify the intron/*sigL* junction (Table S4). Loss of SigL expression was verified by RT-PCR using primers AC106 *sigL*-F and AC107 *sigL*-R (Figure S1), and *sigL* disruption mutants were cured of the Clostron plasmid by serial passage on BHIS without selection.

### Generation of complemented and *sigL-*overexpressing strains

The pMTL8000-series of shuttle vectors was used for construction of *sigL* complementation and overexpression plasmids. Primers AC55 *sigL*-F and AC56 *sigL*-R were designed to amplify the coding sequence approximately 300 base pairs upstream of the *sigL* gene. The resulting PCR products were cleaned and digested with restriction enzymes allowing them to be cloned into either pMTL82151 and expressed under the control of their endogenous promotor (complementation) or pMTL82153 and expressed under the control of a constitutive promotor (Pfdx, overexpression). The resulting plasmids were transformed into CA434 and conjugated into *C. difficile sigL* mutants. Successfully complemented strains were selected on BHIS containing thiamphenicol.

### In silico analysis of SigL-dependent promoters

Twelve putative promoter sequences for genes or operons with previously established downregulation in *sigL* mutants (Soutourina et al., 2020) were used to generate a positional weight matrix (PWM). Genomes of 630 (NCBI Assembly GCA_000009205.2), BI1 (GCA_000211235.1) and a RT078 strain, QCD-23m63 (GCA_000155065.1), were searched for promoter sites based on the *C. difficile sigL* PWM using FIMO (Find Individual Motif Occurrences) (Grant et al., 2011). Identified *sigL* sites in 630, BI1 and RT078 with FIMO q-values less than 0.05 located less than 200 bp upstream of the gene and/or operon were compiled, tabulated and cross-referenced to previous *C. difficile* 630 *sigL* promoters, enhancer binding protein (EBP) upstream activator sequences (UAS), and *sigL::erm* microarray results (Nie et al., 2019;Soutourina et al., 2020). *C. difficile* 630 operons were determined using Prokaryotic Operon DataBase (ProOpDB) (Taboada et al., 2012) were used. The sequence and position of EBP UAS identified by Nie, et. al., in BI1 and RT078 genomes were determined using BLASTn.

### Protein processing and iTRAQ labeling

*C. difficile* was grown to an OD_600_ of either 0.6 (mid-exponential), 0.8 (late-exponential) or 1.0 (stationary) in BHI. Cell pellets were resuspended in 400 μL of 0.5 M triethylammonium bicarbonate (TEAB) buffer and sonicated (five pulses of 15 seconds duration each) at an amplitude setting of 4 for lysis. Lysates were centrifuged at 20,000xg for 30 minutes at 4°C to clear debris. Total recovered proteins were quantitated using a BCA Protein Assay (Pierce, Rockford, IL).

A total of 100 μg of protein in 20 μL of 0.5 M TEAB from each sample was denatured (1 μL of 2% SDS), reduced (1 μL of 100 mM tris-(2-carboxyethyl) phosphine), and alkylated (1 μL of 84 mM iodoacetamide). Trypsin was added at a ratio of 1:10 and the digestion reaction was incubated for 18 hours at 48°C. iTRAQ reagent labeling was performed according to the 8-plex iTRAQ kit instructions (AB SCIEX, Framingham, MA).

### 2D-LC fractionation

Strong cationic exchange (SCX) fractionation was performed on a passivated Waters 600E HPLC system, using a 4.6 × 250 mm polysulfoethyl aspartamide column (PolyLC, Maryland, U.S.A.) at a flow rate of 1 mL/min. Fifteen SCX fractions were collected and dried down completely, then resuspended in 9 μl of 2% (v/v) acetonitrile (ACN), 0.1% (v/v) trifluoroacetic acid (TFA).

For the second-dimension fractionation by reverse phase C18 nanoflow-LC, each SCX fraction was autoinjected onto a Chromolith CapRod column (150 × 0.1 mm, Merck) using a 5 μL injector loop on a Tempo LC MALDI Spotting system (ABI-MDS/Sciex). Separation was over a 50 min solvent gradient from 2% ACN and 0.1% TFA (v/v) to 80% ACN and 0.1% TFA (v/v) with a flow rate of 2.5 μL/min. An equal flow of MALDI matrix solution was added post-column (7 mg/mL recrystallized CHCA (α-cyano-hydroxycinnamic acid), 2 mg/mL ammonium phosphate, 0.1% TFA, 80% ACN). The combined eluent was automatically spotted onto a stainless steel MALDI target plate every 6 seconds (0.6 μl per spot), for a total of 370 spots per original SCX fraction.

### Mass spectrometry analysis and protein quantitation

MALDI target plates were analyzed in a data-dependent manner on an ABSciexTripleTOF 5600+ mass spectrometer (AB SCIEX, Framingham, MA). MS spectra were acquired from each sample spot using 500 laser shots per spot, laser intensity 3200. The highest peak of each observed m/z value was selected for subsequent MS/MS analysis.

Up to 2500 laser shots at laser power 4200 were accumulated for each MS/MS spectrum. Protein identification and quantitation was performed using the Paragon algorithm implemented in Protein Pilot 3.0 software by searching the acquired MS and MS/MS spectra from all 15 plates against the *C. difficile* strain BI-1 protein database, or the 078 representative strain QCD-23m63 database, plus common contaminants. The Protein Pilot Unused score cutoff of > 0.82 (1% global false discovery rate) was calculated from the slope of the accumulated Decoy database hits by the Proteomics System Performance Evaluation Pipeline (PSPEP) program [72]. Proteins with at least one peptide greater than 95% confidence (score based on the number of matches between the data and the theoretical fragment ions) and a Protein Pilot Unused score of > 0.82 were considered as valid identifications (IDs).

### Identification of differentially abundant proteins

A pooled sample consisting of equal amounts of each protein sample was used as a technical replicate. iTRAQ tags 113 and 121 were used to tag equal amounts of the same pooled sample. A receiver operating characteristic (ROC) curve was plotted using the technical replicate ratios, 119:121 and 121:119, as true negatives and all studied ratios as true positives. A 2-fold change cut-off was set to define proteins altered in abundance between wt and respective *sigL::erm* mutants.

### PSII extraction and detection

PSII extraction and quantitation was performed by previously published methods (Chu et al., 2016) with minor modifications. *C. difficile* vegetative cells were grown to an OD_600_=1.2 in BHI at 37°C. Cells were pelleted by centrifugation for 10 minutes at 4000 rpm, and the supernatant was discarded. Pellets were resuspended in 0.1M EDTA-triethanolamine buffer pH 7.0 for 20 minutes, and subsequently centrifuged for 10 minutes at 4000 rpm. The supernatant was collected, serially diluted, and dot-blotted onto activated PVDF membranes, and probed with PSII-specific antiserum. Antibodies were detected using anti-Rabbit HRP secondary and FemtoWest Maximum sensitivity substrate (Pierce).

### Autolysis assays

Autolysis of wild type and *sigL* mutant strains was measured using the method described previously (El Meouche et al., 2013). 50 mL *C. difficile* cultures were grown in BHI to an OD_600_=1.2, and were centrifuged for 10 minutes at 4000 rpm to pellet. The harvested cells were washed twice in PBS and resuspended in 50mL of 50mM phosphate buffer containing 0.01% of Triton X-100. Absorbance values at OD_600_ of resuspended cells were recorded, and cultures were placed back into an anaerobic environment at incubated at 37°C. Time points were harvested every 15 minutes, and autolysis was monitored by measuring absorbance values as a percentage of the initial OD_600_. Each assay was performed in biological triplicate, and photographed at the termination of the experiment.

### Autoaggregation analysis

Autoaggregation was measured by the method of Faulds-Pain et. al (Faulds-Pain et al., 2014) with modifications. Overnight cultures of *C. difficile* wild type and *sigL* mutants were diluted 1:50 in BHI, and 5mL of culture was separated into pre-reduced glass culture tubes and incubated upright at 37°C. Each hour, six culture tubes were removed from incubation and the OD_600_ was determined. The OD_600_ was directly read from the top 1mL of culture in three “un-mixed” fractions per time point, and the remaining three tubes were vortexed and the 0D_600_ was measured constituting “mixed” fractions. Aggregated cells settle from suspension and are not recorded in the OD_600_ from “unmixed” fractions but are detected in the “mixed” fractions. The percent agglutination of aggregative *C. difficile* vegetative cells was calculated as the difference between the OD_600_ of the whole (mixed) solution and the OD_600_ of the top 1 ml (unmixed) as a percentage of the whole (mixed) solution OD_600_.

### Biofilm assays

The influence of SigL expression on *C. difficile* biofilm formation was evaluated using a biofilm assay as previously reported (Pantaléon et al., 2015) with modifications. Overnight *C. difficile* cultures were diluted 1:50 in BHI and distributed into 12-well plates, 2mL per well. Plates were wrapped to prevent evaporation and incubated anaerobically at 37°C for 72 hours. Following incubation, the culture supernatant was removed and the biofilms were washed twice with PBS and then incubated at 37°C with .02% crystal violet to fix and stain. The crystal violet was subsequently removed, and the biofilms were washed an additional two times with PBS and photographed. To measure biofilm formation, crystal violet retained by the biomass was released with acetone/ethanol and quantitated by taking absorbance measurements at 570nm in technical quadruplicate for three individual wells.

### *C. difficile* adhesion assays to Caco2-BBE human intestinal epithelial cells

*C. difficile* attachment to human colonic epithelial cells was performed using the modified methods of Merrigan et al (Merrigan et al., 2013). C2BBe1 cells were obtained from the American Type Culture Collection (ATCC) and were grown in 2mL DMEM containing 5% fetal bovine serum to confluent monolayers in 6 well tissue culture plates. Once confluent, the monolayers were incubated overnight in 2mL DMEM without serum. *C. difficile* strains to be analyzed were subcultured from overnight growth in BHI 1:50 into BHI and grown to an OD_600_=0.4. C2BBe1 cells were enumerated and multiplicity of infection (MOI) was calculated for each individual strain tested with a target of MOI=50. *C. difficile* was harvested at OD_600_=0.4 and centrifuged 5 minutes at 4000 rpm to remove BHI. *C. difficile* pellets and confluent monolayers were brought into the anaerobic chamber. *C. difficile* pellets were resuspended in DMEM without serum and placed on the monolayers to adhere for a total of 40 minutes at 37°C anaerobically. The inoculum was serially diluted and titered on BHIS with antibiotics as appropriate to calculate MOI. Following attachment, the supernatant was removed, and the monolayers were washed twice with PBS to remove non-adherent bacteria. Monolayers were resuspended in 1mL PBS and scraped up to quantify attachment. Recovered *C. difficile* were serially diluted in PBS and plated on BHIS with antibiotics as appropriate to enumerate adherent levels of bacteria.

### Sporulation assessments

To determine the kinetics of CD sporulation, overnight cultures of CD in BHIS were subcultured 1:50 into BHI and grown to an OD_600_=0.4. These cultures were used to inoculate 50mL of BHI per time point analyzed. At each time point, cultures were harvested by centrifugation for 10 minutes at 4000 rpm, and the supernatant was removed. Cell pellets were washed once in 1mL PBS, centrifuged, and resuspended in 1mL PBS prior to heat shocked for 20 minutes at 60°C. Following heat shock to inactivate remaining vegetative cells, the remaining spores were serially diluted in PBS and plated on BHIS-Taurocholate plates. Plates were incubated anaerobically for 24 hours at 37°C and enumerated.

### Motility assays

Quantitative assessment of C. *difficile* motility was performed as previously described (Baban et al., 2013). Briefly, overnight cultures of *C. difficile* grown in BHIS were diluted 1:50 in fresh BHI and grown to mid-exponential growth phase (OD_600_nm=0.6) at 37°C in anaerobic conditions. 5μL of culture was then stab inoculated into the center of one well of a 6-well tissue culture plate filled with 20mL of *C. difficile* motility agar (BHI+0.3% agar). Culture plates were wrapped in plastic wrap, placed into a container with damp paper towels to prevent evaporation, and grown anaerobically for 72 hours at 37°C. Following incubation, the diameter of the swim radius was measured and averaged. Results represent the average of three independent experiments.

### Secreted toxin analyses

Secreted *C. difficile* toxin was detected by immunoblotting using monoclonal antibodies against both TcdA and TcdB. *C. difficile* cultures were grown in 35mL quantities for times as indicated, and vegetative cells were removed by centrifugation at 4 °C for 10 minutes at 4000 rpm. 30mL of culture supernatant was harvested and concentrated 1:50 on a 100kDa molecular weight cut-off column (EMD Millipore, Billerica, MA). Concentrated proteins were quantitated using a standard BCA assay (ThermoFisher, Waltham, MA), and 50ug of protein was loaded onto 7.5% TGX acrylamide gels (BioRad, Hercules, CA) for electrophoresis. Proteins were transferred to nitrocellulose membranes (BioRad) and TcdA and TcdB were immunoblotted using specific monoclonal antisera (Abcam) at 1:5000 dilution. Samples were detected using a secondary horseradish-peroxidase conjugated rabbit anti-mouse IgG antibody (BioRad) at 1:10,000 concentration. Antibodies were detected using FemtoWest Max Sensitivity substrate (ThermoFisher).

The Alere Wampole A/B Toxin ELISA kit (Alere, Atlanta GA) was utilized to determine relative levels of combined TcdA and TcdB secreted by *C. difficile* strains. Strains were grown for 72 hours in BHI broth, and supernatant was harvested by centrifugation. Supernatants were diluted 1:10, and 1:100 in fresh BHI and processed for ELISA as per the manufacturer’s instructions to determine the appropriate working concentration. Prior to analysis, the amount of total protein was normalized using the BCA assay (Pierce) to allow direct comparison of toxin secretion levels among different *C. difficile* strains. Following assay processing as per the manufacturer’s instructions, quantification of *C. difficile* toxin was performed by reading absorbance at 450nm using an automated microtiter plate reader (Bio-Tek, Winooski VT).

### Statistical Methods

The Excel-Stat application was used for statistical analysis. Student’s t-tests was used compute differences between two groups, and errors bars calculated from standard deviation(s). Analysis of Variance (ANOVA) was used to compare >2 groups, followed by Tukey’s HSD test for post-hoc analysis.

## Acknowledgements

This work was supported by funding from the National Institutes of Health [R33AI121590531(GV)] and the US Dept. of Veterans Affairs [Research Career Scientist Award IK6BX003789(GV); and Merit Award I01BX001183(GV)]. The authors acknowledge Aaron Brussels for expert technical assistance, the University of Arizona Mass Spectrometry Consortium for mass spectrometry studies and data compilation, and members of the Vedantam and Viswanathan laboratories for helpful discussions and critical reading of the manuscript.

