## Supplementary figures and images for "The Alternative Sigma Factor SigL Influences *Clostridioides difficile* Toxin Production, Sporulation, and Cell Surface Properties"

### Supplemental Figure 1

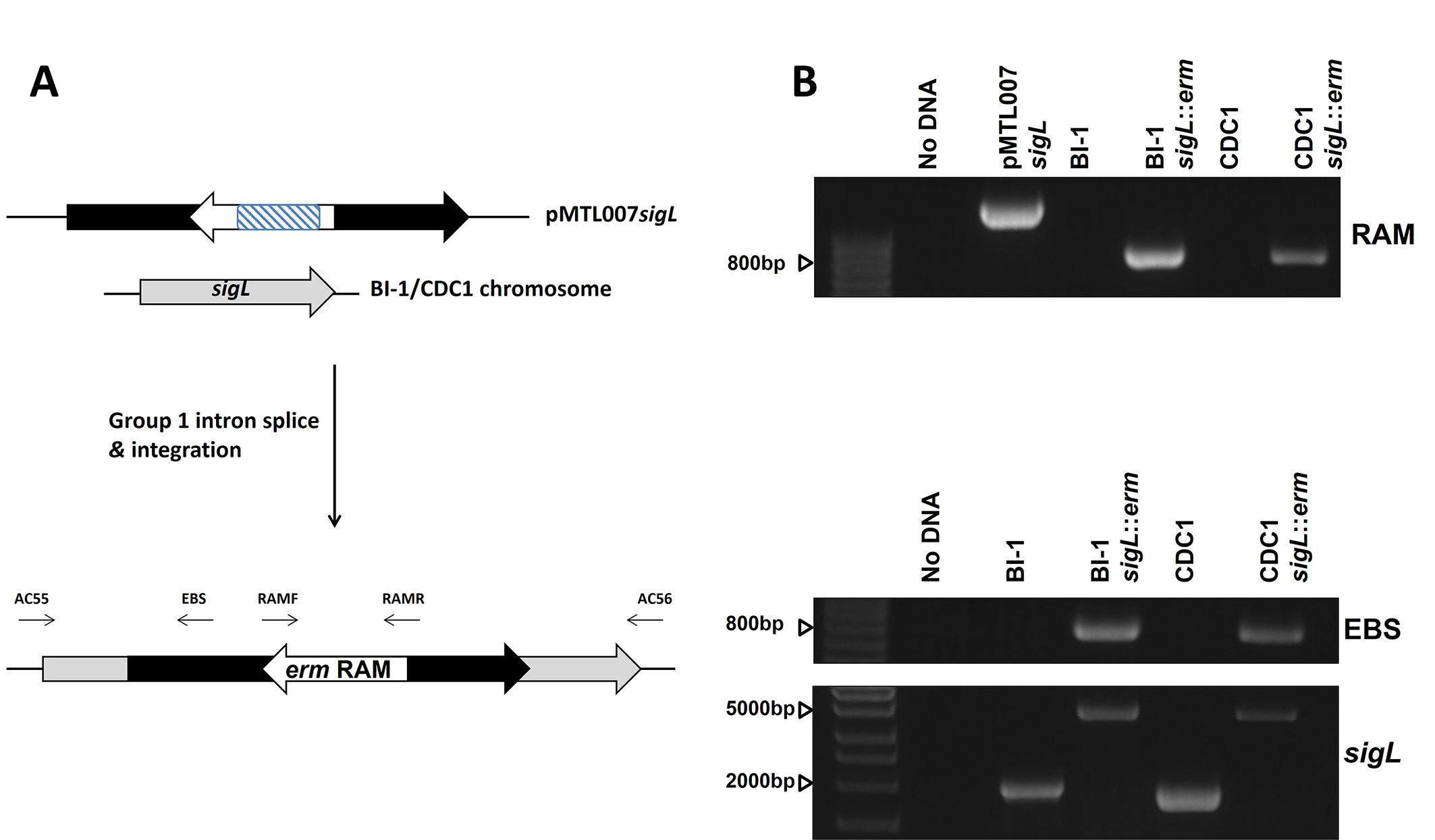

### Supplemental Figure 2

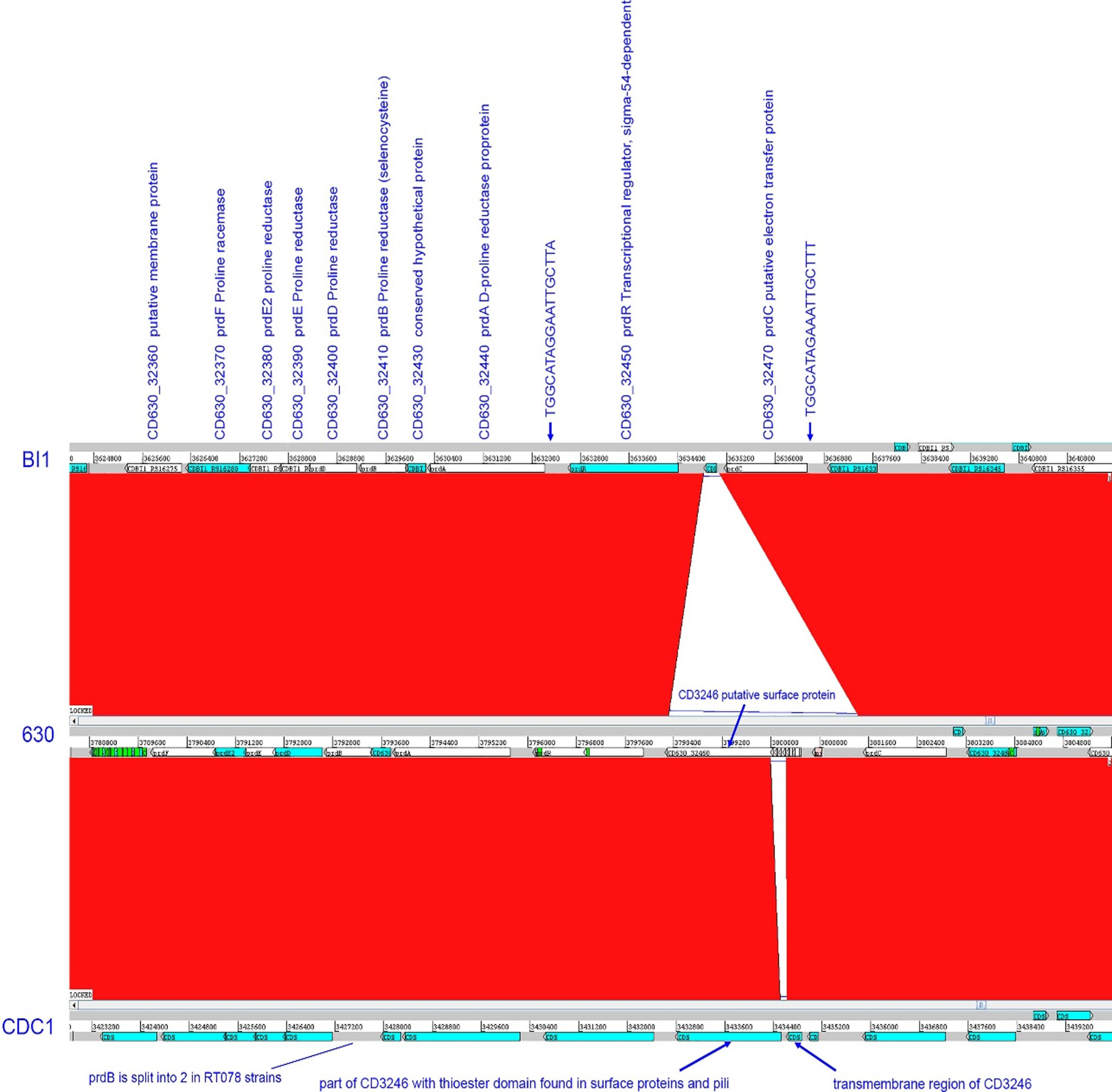

### Supplemental Figure 3

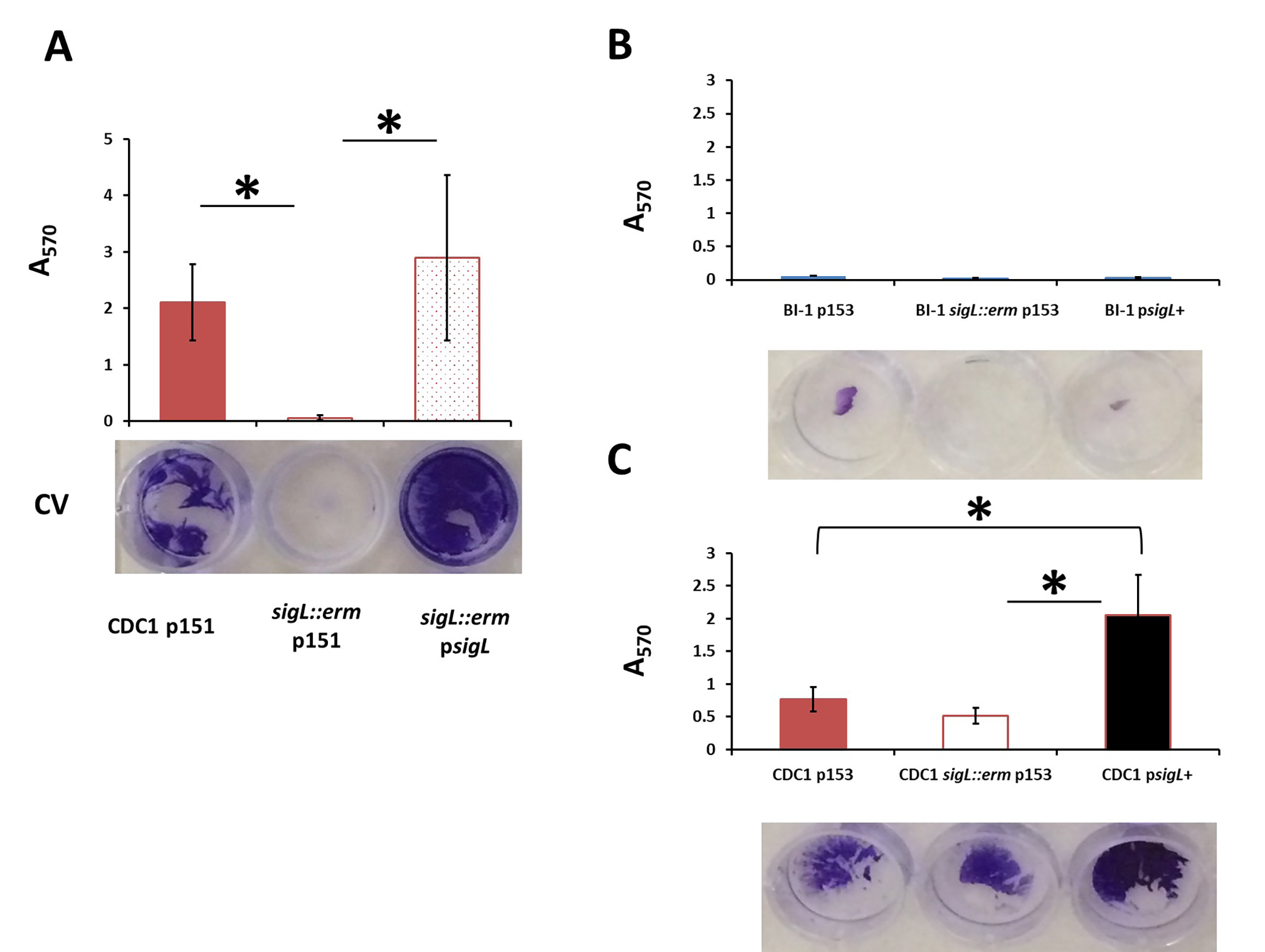

### Supplemental Figure 4

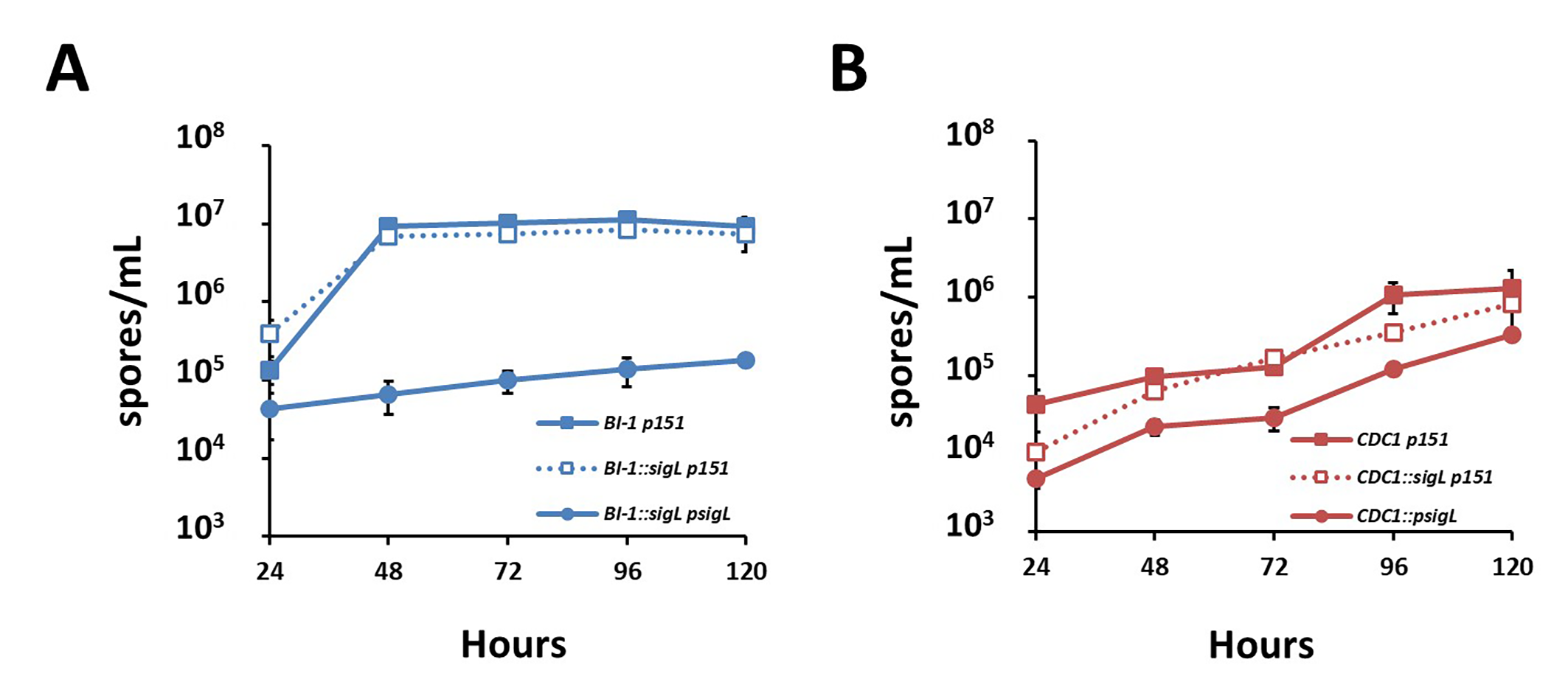

### Supplemental Figure 5

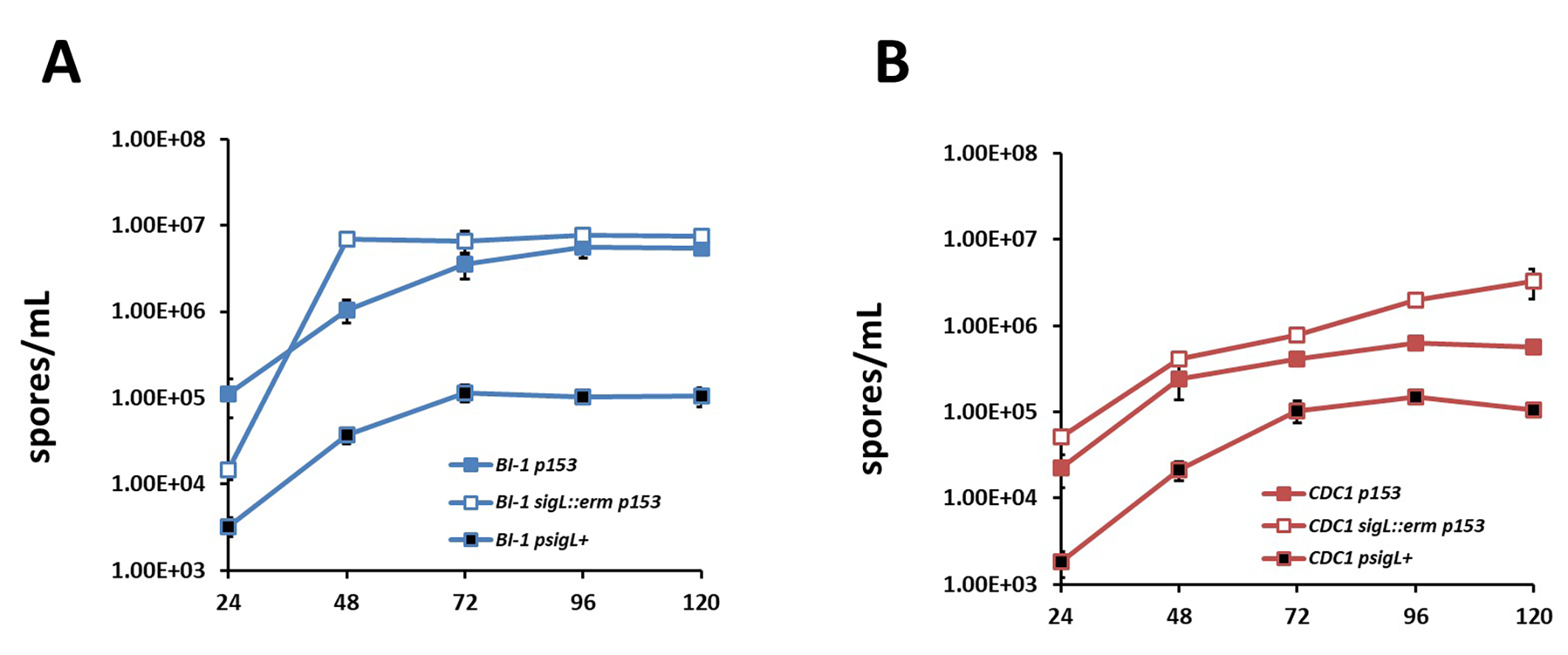

### Supplemental Figure 6

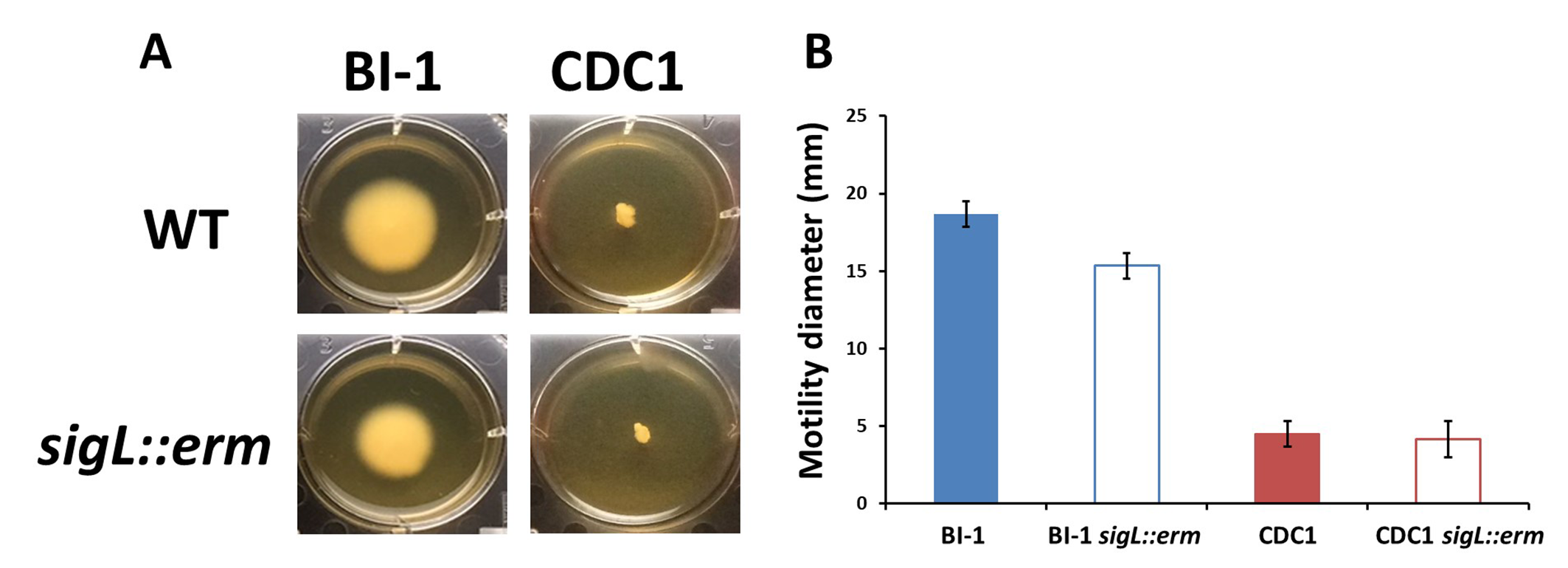

### Supplemental Figure 7

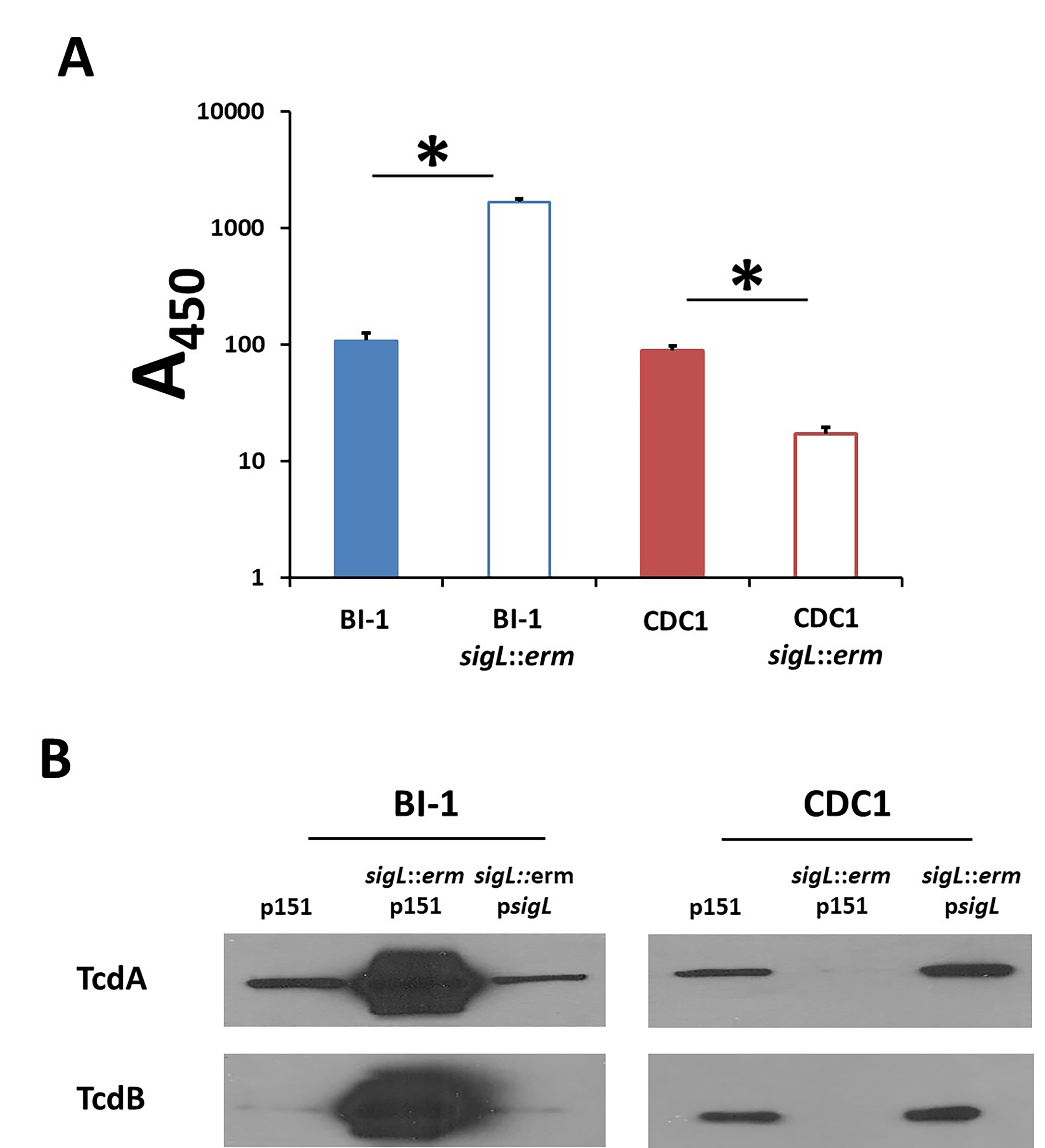
