## Supplemental Tables 3 and 4 for "The Alternative Sigma Factor SigL Influences *Clostridioides difficile* Toxin Production, Sporulation, and Cell Surface Properties"

### Slide 1
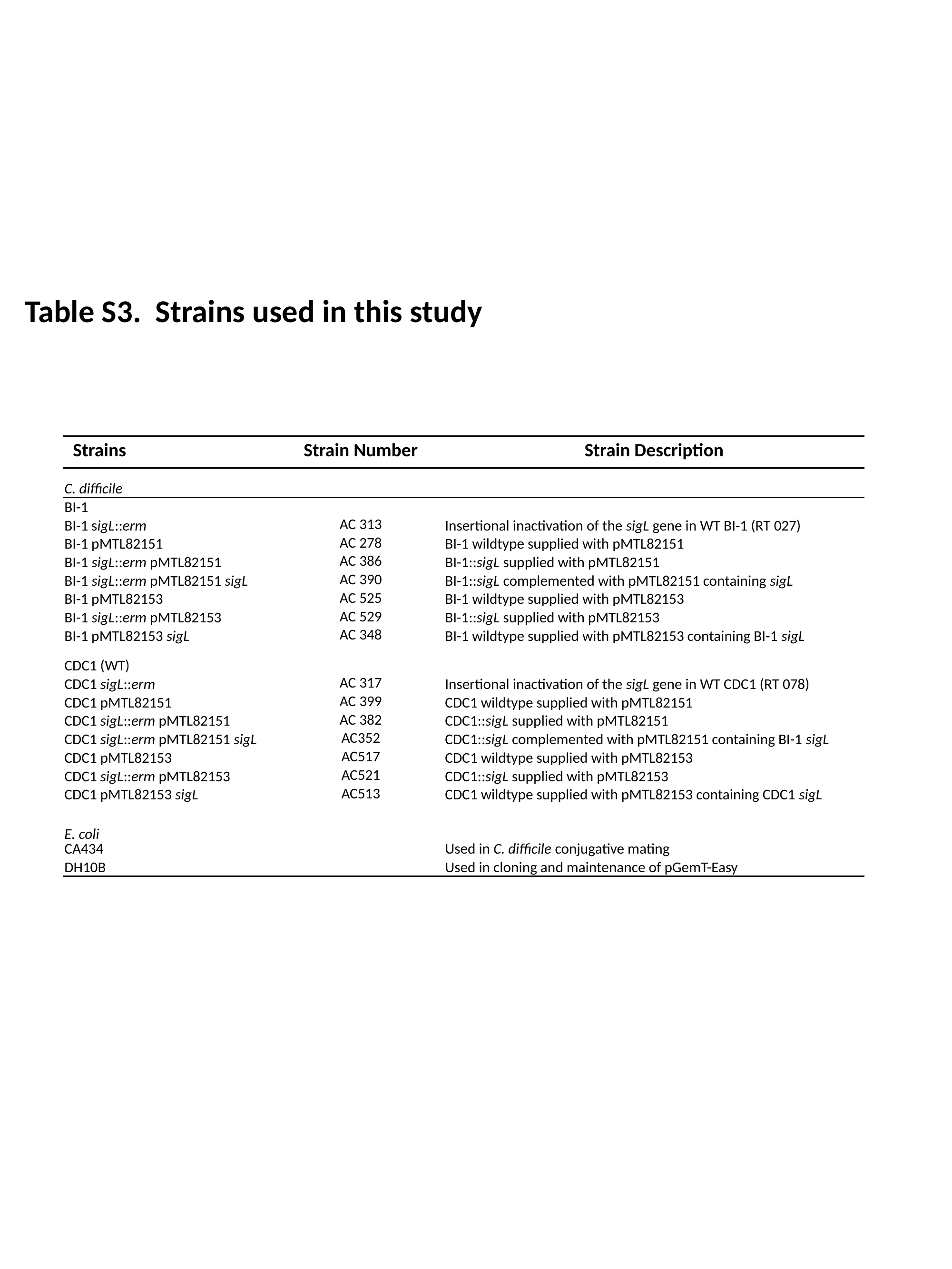

Table S3. Strains used in this study
| Strains | Strain Number | Strain Description |
| --- | --- | --- |
| C. difficile | | |
| BI-1 | | |
| BI-1 sigL::erm | AC 313 | Insertional inactivation of the sigL gene in WT BI-1 (RT 027) |
| BI-1 pMTL82151 | AC 278 | BI-1 wildtype supplied with pMTL82151 |
| BI-1 sigL::erm pMTL82151 | AC 386 | BI-1::sigL supplied with pMTL82151 |
| BI-1 sigL::erm pMTL82151 sigL | AC 390 | BI-1::sigL complemented with pMTL82151 containing sigL |
| BI-1 pMTL82153 | AC 525 | BI-1 wildtype supplied with pMTL82153 |
| BI-1 sigL::erm pMTL82153 | AC 529 | BI-1::sigL supplied with pMTL82153 |
| BI-1 pMTL82153 sigL | AC 348 | BI-1 wildtype supplied with pMTL82153 containing BI-1 sigL |
| CDC1 (WT) | | |
| CDC1 sigL::erm | AC 317 | Insertional inactivation of the sigL gene in WT CDC1 (RT 078) |
| CDC1 pMTL82151 | AC 399 | CDC1 wildtype supplied with pMTL82151 |
| CDC1 sigL::erm pMTL82151 | AC 382 | CDC1::sigL supplied with pMTL82151 |
| CDC1 sigL::erm pMTL82151 sigL | AC352 | CDC1::sigL complemented with pMTL82151 containing BI-1 sigL |
| CDC1 pMTL82153 | AC517 | CDC1 wildtype supplied with pMTL82153 |
| CDC1 sigL::erm pMTL82153 | AC521 | CDC1::sigL supplied with pMTL82153 |
| CDC1 pMTL82153 sigL | AC513 | CDC1 wildtype supplied with pMTL82153 containing CDC1 sigL |
| E. coli CA434 | | Used in C. difficile conjugative mating |
| DH10B | | Used in cloning and maintenance of pGemT-Easy |

### Slide 2
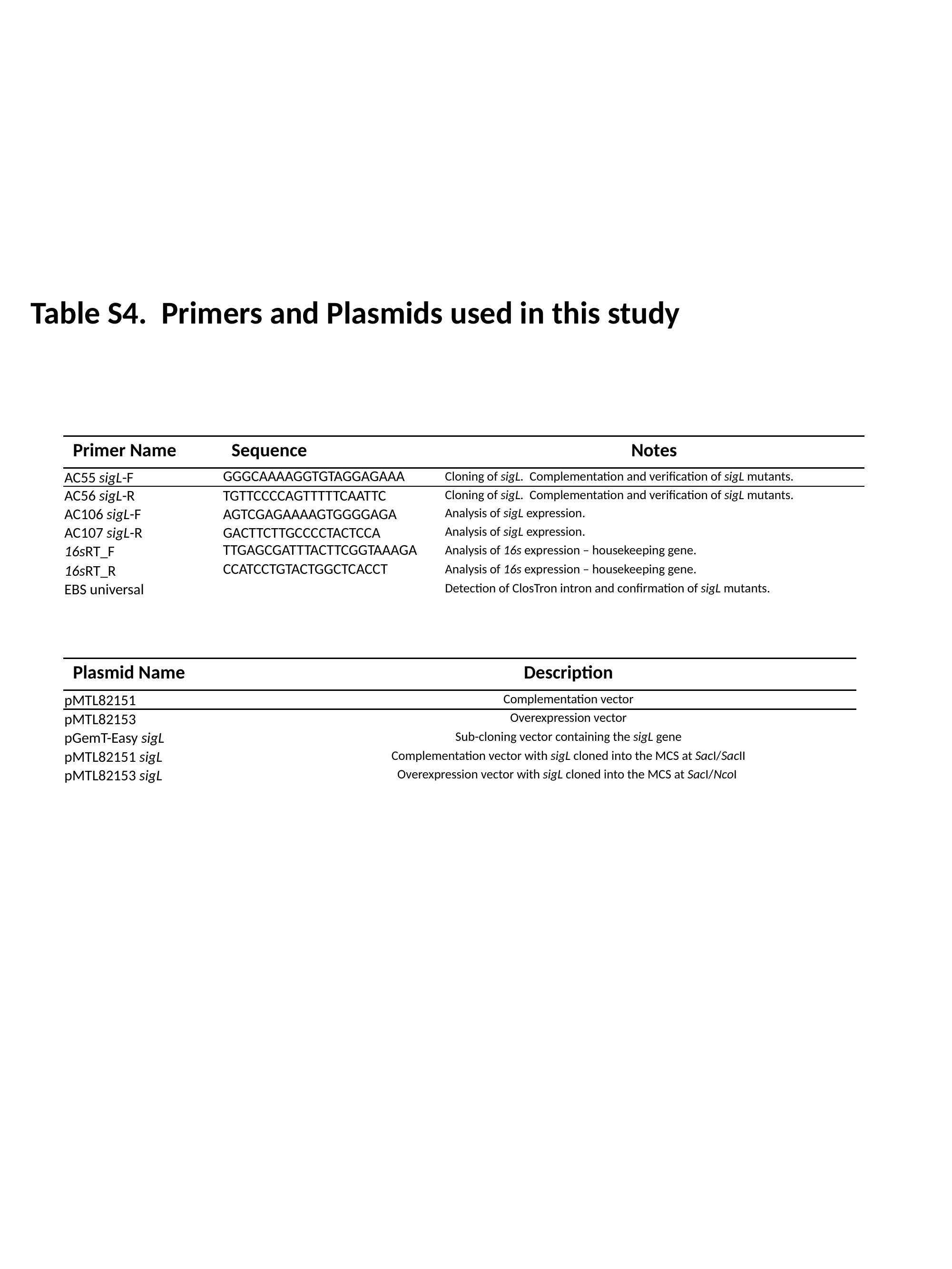

Table S4. Primers and Plasmids used in this study
| Primer Name | Sequence | Notes |
| --- | --- | --- |
| AC55 sigL-F | GGGCAAAAGGTGTAGGAGAAA | Cloning of sigL. Complementation and verification of sigL mutants. |
| AC56 sigL-R | tgttccccagtttttcaattc | Cloning of sigL. Complementation and verification of sigL mutants. |
| AC106 sigL-F | agtcgagaaaagtggggaga | Analysis of sigL expression. |
| AC107 sigL-R | gacttcttgcccctactcca | Analysis of sigL expression. |
| 16sRT\_F | TTGAGCGATTTACTTCGGTAAAGA | Analysis of 16s expression – housekeeping gene. |
| 16sRT\_R | CCATCCTGTACTGGCTCACCT | Analysis of 16s expression – housekeeping gene. |
| EBS universal | | Detection of ClosTron intron and confirmation of sigL mutants. |
| Plasmid Name | Description |
| --- | --- |
| pMTL82151 | Complementation vector |
| pMTL82153 | Overexpression vector |
| pGemT-Easy sigL | Sub-cloning vector containing the sigL gene |
| pMTL82151 sigL | Complementation vector with sigL cloned into the MCS at SacI/SacII |
| pMTL82153 sigL | Overexpression vector with sigL cloned into the MCS at SacI/NcoI |
