## Supplementary Figure Captions for "The Alternative Sigma Factor SigL Influences *Clostridioides difficile* Toxin Production, Sporulation, and Cell Surface Properties"

**Figure S1. *sigL* inactivation by Clostron mutagenesis (A).** Schematic showing Clostron mutagenesis with the targeting plasmid pMTL007sigL (adapted from Underwood et al, 2009). The disrupted locus in the mutant, with primer binding sites, is also shown. **(B).** Confirmation of *sigL* inactivation in BI-1 and CDC1 strains. The three panels show PCR products generated with the indicated templates using primers RAMF + RAMR (confirmation of intron integration), AC55 + EBS (site-specific confirmation of integration site), and AC55 + AC56 (size shift in disrupted *sigL*) respectively.

**Table S1**. ***sigL* motif identity with FIMO scores in *C. difficile* strains 630, BI-1 and CDC1.** 30 *sigL*-dependent operons identified by Soutorina et al are listed in the first column for comparison. CD630_34780 was identified in our study and not in Soutourina et al. Associated strains, loci, names, FIMO scores/q-values and motif sequences are listed for the identified motifs. Two loci of the 30 previously identified are not identified in CDC1. Lower FIMO scores are bolded, and atypical nucleotides in the motif sequences are shown in red.

**Figure S2. Comparison of *prd* locus in 630, BI-1, and CDC1.**  The genomes of *C. difficile* BI-1, CDC1, and 630 were compared using BLASTn, and Artemis comparison tool (ACT) was used to generate the pairwise comparisons. CD3246, encoding a putative surface protein, is present between *prdC* and *prdR* in 630. This segment is largely truncated in CDC1, and replaced by a gene encoding a putative transposase in BI-1.

**Table S2. Growth and *sigL*-dependent proteomic changes in BI-1 and CDC1.** Total protein abundances of BI-1, CDC1, and their respective *sigL::erm* derivatives were compared using iTRAQ labeling at early-exponential, late-exponential, and stationary growth phases. A 2-fold cutoff was used to determine significant change in protein abundance.

**Figure S3. Complementation of CDC1 *sigL::erm* restores the ability to form biofilm.** WT CDC1 was conjugated with empty vector pMTL82151 (p151). *sigL::erm* was conjugated with empty vector or p151 expressing *sigL* under an endogenous promoter (p*sigL*). Biofilm was quantified by the release of crystal violet retained by the biomass after 72 hours, using acetone/ethanol. Absorbance was measured at 570 nm. Samples were grown in biological triplicate and compared using ANOVA and Tukey’s HSD post-hoc test. *p<.05

**Figure S4. Plasmid complementation of *sigL* results in fewer spores.** WT *C. difficile* strains were conjugated with empty vector pMTL82151 (p151). *sigL::erm* in BI-1 and CDC1 backgrounds were conjugated with empty vector or p151 expressing sigL under an endogenous promoter (p*sigL*). Spores were enumerated by heat shocking and plating cultures on BHIS-Taurocholate at each time point. **(A).** Spore enumeration over 120 hours in BI-1 and BI-1 sigL::erm strains **(B).** Spore enumeration over 120 hours in CDC1 and CDC1 sigL::erm strains. Cultures were grown in biological triplicate.

**Figure S5. *sigL* overexpression reduces sporulation in BI-1 and CDC1.** WT *C. difficile* strains were conjugated with empty vector pMTL82153 (p153). *sigL::erm* in BI-1 and CDC1 backgrounds were conjugated with empty vector or p153 expressing sigL under a constitutive promoter (p*sigL+*). Spores were enumerated by heat shocking and plating cultures on BHIS-Taurocholate at each time point. **(A).** Spore enumeration over 120 hours in BI-1 strains **(B).** Spore enumeration over 120 hours in CDC1 strains. Cultures were grown in biological triplicate.

**Figure S6. Loss of *sigL* expression does not impact flagellar motility in BI-1.** Motility was measured as the diameter of the swim radius in soft agar. CDC1 is aflagellate and was used as a negative control for motility. **(A).** Representative images of swim radius for *C. difficile* strains **(B).** Swim radius of BI-1 and CDC-1 strains with respective *sigL::erm* strains. Assays were conducted in biological triplicate and compared using ANOVA.

**Figure S7. Verification and complementation of *sigL* toxin phenotypes. (A).** Total toxin (TcdA and TcdB) was measured by ELISA in *C. difficile* strains after 72 hours. The absorbance values at 450 nm represent an average of four samples. *C. difficile* strains and respective *sigL::erm* strains were compared using ANOVA. *p<.05 **(B).** BI-1 and CDC1 strains, and respective *sigL::erm* mutants were conjugated with empty complementation vector p151. *sigL::erm* strains were conjugated with p151 expressing *sigL* (p*sigL*). WT, mutant, and complemented strains were immunoblotted for TcdA and TcdB.

**Table S3.** Strains used in this study.

**Table S4.** Primers and plasmids used in this study.
